# In vivo kinetics of protein degradation by individual proteasomes

**DOI:** 10.64898/2026.01.19.700426

**Authors:** Maximilian F. Madern, David Haselbach, Marvin E. Tanenbaum

## Abstract

Protein degradation by the proteasome is central to cellular homeostasis and has been studied extensively using biochemical and structural studies. Despite an in-depth understanding of core proteolytic activity, it has remained largely unresolved how individual proteasomes process substrates inside living cells where many substrate types and co-factors exist. Here, we establish a live-cell single-molecule imaging approach that enables direct visualization and quantification of protein degradation by individual proteasomes. Using this approach, we find that substrate identity, folding and protein-protein interaction have a surprisingly modest impact on processing efficiency, whereas the mode of substrate engagement greatly impacts substrate processing; degradation initiated from protein termini typically proceeds rapidly and with high processivity, whereas internal engagement constitutes a distinct processing mode that exhibits poor processivity and a specific requirement for the AAA+ family ATPase p97/VCP. Furthermore, degradation initiated from opposite termini proceeds with asymmetric rates in a sequence-dependent manner, demonstrating that directionality is an important feature of proteasomal processing *in vivo*. Notably, poly-glutamine substrates associated with neurodegenerative disease are efficiently degraded from one terminus but resist degradation when engaged from the opposite terminus, highlighting the importance of substrate engagement mode. Together, our results show that different modes of substrate engagement lead to different proteasomal processing outcomes *in vivo* and revise the prevailing view of the proteasome as a uniform degradation machine.

## Introduction

Protein degradation is essential for cellular homeostasis and precise control of protein expression levels. The controlled degradation of proteins plays critical roles in processes such as development, the cell cycle, and signal transduction. Conversely, perturbations in degradation pathways are associated with diverse human pathologies, most notably neurodegenerative disorders, in which a failure to degrade protein aggregates is associated with severe toxicity^1-4^.

In eukaryotes, the ubiquitin–proteasome system (UPS) constitutes the major pathway for protein degradation. The proteasome is a 2.5 MDa macromolecular machine that recognizes and degrades polyubiquitinated proteins. Proteasomal degradation typically begins with interactions of ubiquitin adapters on the proteasome with polyubiquitin chains on substrates^5,6,7^. Productive degradation subsequently requires engagement of an unstructured region of the substrate by the proteasome’s internal AAA+ subunits^8,9^. Upon engagement, substrates are unfolded and translocated into the proteolytic core by ATP-driven motors, coupled to substrate deubiquitination^10,11^. Within the core particle, the polypeptide is cleaved into small peptides, which are released for reuse of their constituent amino acids^12,13^. Interestingly, *in vitro* work has revealed that the proteasome can engage with substrates through different modes, initiating decay from the N- or C-terminus, or through substrate-internal engagement, highlighting the complexity of protein decay^8,14^. Proteasome activity is further regulated by an extensive network of cofactors and adaptors, including ubiquitin ligases, deubiquitinating enzymes, substrate-shuttling factors and ATP-dependent unfoldases such as p97/VCP, which together ensure that the appropriate proteins are targeted for degradation^15-17^.

While *in vitro* studies have provided important insights into the core proteolytic activity, the limited substrate diversity and the absence of the large set of regulatory factors in such *in vitro* assays has limited analysis of protein degradation in its full complexity. *In vivo* methods, for example using fluorescently-tagged substrates, can track bulk substrate decay over time^18-21^, but such methods do not provide the molecular resolution required to study substrate engagement and processing in detail. Specifically, these assays do not report on the site of substrate engagement, the rate of substrate recruitment, nor the rate, processivity or directionality of degradation. As such, a precise molecular understanding of substrate engagement and processing *in vivo* is currently lacking.

In this study, we overcome these challenges by developing a molecular-resolution *in vivo* imaging approach that enables visualization and precise quantitation of processing of individual substrates by single proteasomes. Using this assay, we unravel how substrate diversity and engagement mode impact substrate processing by the proteasome, revealing mechanistic principles governing *in vivo* protein degradation, and establishing single-molecule protein decay imaging as a powerful tool to understand proteostasis.

## Results

### A live-cell assay to visualize degradation by single proteasomes

In order to visualize degradation of individual proteasome substrates in living cells, we sought to engineer very long protein substrates with high and uniform fluorescent labeling along the protein. We reasoned that degradation of a long fluorescent substrate would result in a gradual decrease in fluorescence over time, which could be tracked by single-molecule microscopy and would report on proteasome degradation kinetics (Fig. 1a). Importantly, this approach would allow disentanglement of distinct sub-steps of protein degradation, including initiation of decay (i.e. substrate recruitment) and processive substrate degradation. To create a long, brightly-labeled polypeptide, we made use of stopless-ORF circRNA (socRNA) technology that we recently developed, which employs rolling-circle translation to generate very long, fluorescently labeled proteins^22^. We refer to these proteins as socRNA-encoded repeat proteins (SERPs) (Fig. 1a). Fluorescent labeling of SERPs was achieved using the SunTag system^23^, in which peptide epitopes are bound by a fluorescently labeled single-chain antibody that is constitutively expressed in cells, yielding bright, diffraction-limited foci that represent individual SERPs. To facilitate long-term tracking and precise quantitative measurements of individual SERPs, SERPs were tethered to the plasma membrane using the orthologous ALFA-Tag peptide tagging system^24^ (Fig. 1b). To trigger SERP degradation by the proteasome, we co-opted a co-translational ribosome quality control pathway that is activated upon ribosome stalling, and triggers C-terminal SERP ubiquitination followed by SERP release from the ribosome^25,26^. Ribosome stalling was induced by socRNA cleavage through introduction of a let-7a microRNA binding site with perfect sequence complementarity into the socRNA (Fig. 1c, d and Supplementary Fig. 1a). SocRNAs containing the let-7a binding site displayed markedly shortened translation time, consistent with efficient cleavage of socRNAs (Supplementary Fig. 1b). To further facilitate single-molecule protein degradation readouts using this assay, we designed socRNAs that are translated almost exclusively by single ribosomes, so each GFP spot represents a single polypeptide (Supplementary Fig. 1c).

**Fig. 1.**
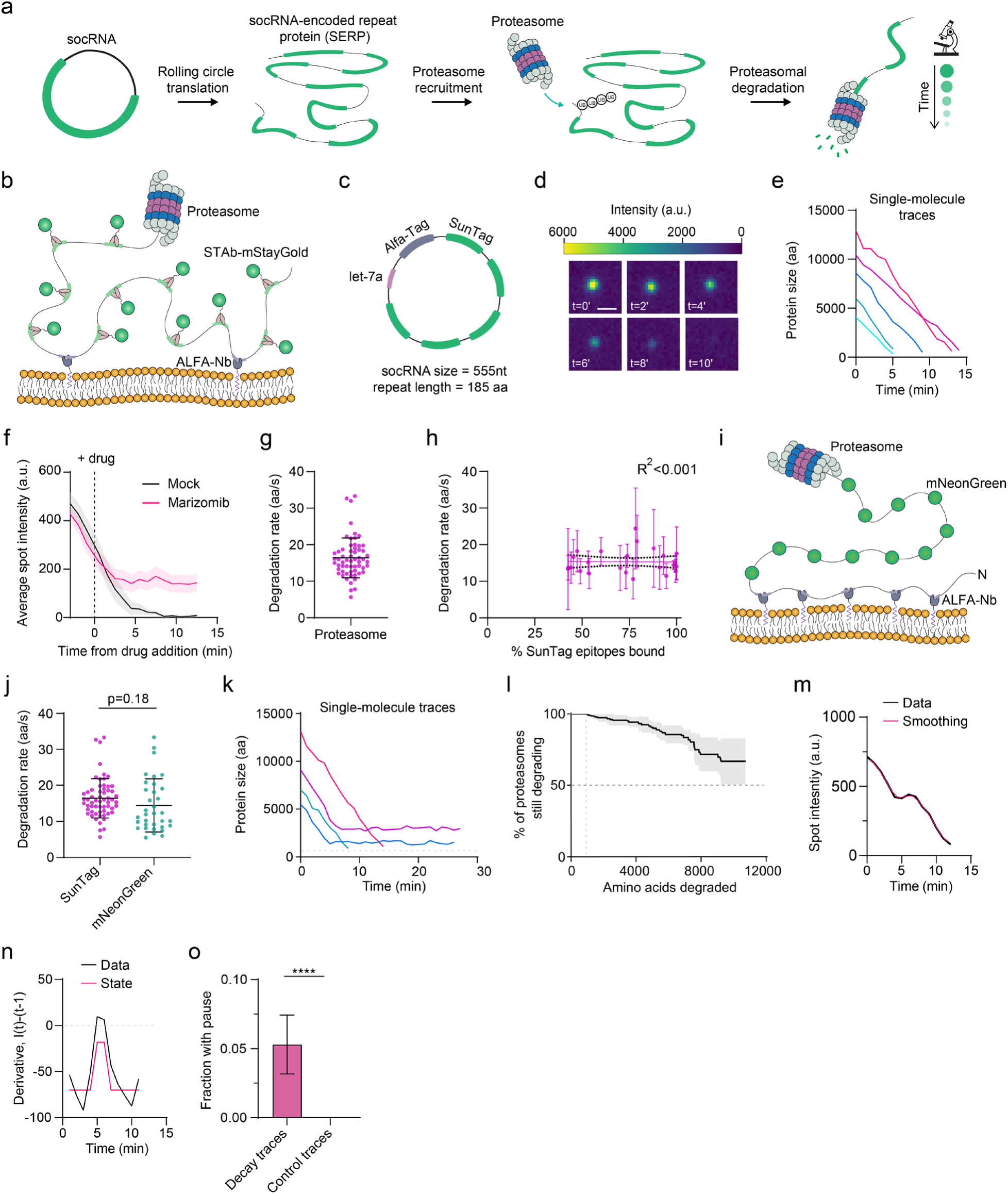
Single-molecule imaging of proteasomal degradation in living cells. **a,** Illustration of the experimental approach to visualize protein degradation by individual proteasomes in real-time. **b,** Schematic of the single-molecule protein degradation imaging assay. Proteasome substrates (SERPs) are fluorescently labeled via SunTag antibodies (STAb) fused to the green fluorescent protein mStayGold, and anchored to the plasma membrane using ALFA-tag nanobodies (ALFA-Nb) fused to a membrane-targeting CAAX motif. **c,** Schematic of a socRNA containing five SunTag epitopes, a single the SunTag ant ALFA-tag epitope, and a sequence fully complementary to the let-7a microRNA, introduced to induce RNA cleavage and proteasomal degradation of the SERP. **d,** Representative example images of a time-lapse movie for a single SERP undergoing degradation. **e,** Representative intensity-time traces of SERP degradation events. Scale bar, 1 μm. **f,** Average SERP fluorescence intensities over time during degradation. Decay events were aligned to the moment of drug addition, indicated by the dashed vertical line. Lines and shaded region indicate mean ± s.d (n = 11 (Mock) and 16 (Marizomib) traces from 2 independent repeats). **g,** Rate of protein degradation by the proteasome. Data points represent individual SERP degradation events. Horizontal line and error bars represent mean ± s.d of medians (n = 60 traces from 3 independent repeats). **h,** Relationship between the stoichiometry of SunTag epitope labeling and decay rate. Each dot and error bars represent the average ± s.d. of the decay rate of SERPs in a single cell. Black line represents fit of the data using a linear regression model, black dashed lines represent 95% CI for the linear regression model (n = 137 traces from 24 different cells). **i,** Schematic of proteasome imaging assay using a mNeonGreen fluorescent protein in the repeated unit of the SERP. Substrates are anchored via an N-terminal 5x ALFA-tag array (see also Supplementary Fig. 1h). **j,** Comparison of decay rates for SERPs encoding the SunTag peptides and mNeonGreen. Horizontal lines and error bars represent mean ± s.d (n = 60 (SunTag) and 34 (mNeonGreen) traces from 3 independent repeats). **k**, Representative intensity-time traces of complete and incomplete degradation traces. Dashed line represents the approximate detection threshold in our assay. **l**, Kaplan-Meier survival curve showing the number of amino acids degraded by single proteasomes before degradation is aborted (i.e. proteasome processivity). Line and shaded region indicate mean ± s.d (n = 170 traces from 5 independent repeats). **m**, Representative intensity-time trace for a SERP degradation event exhibiting a pause. Raw data (black) is overlaid with 2-point moving average (red). **n,** Derivative of the smoothed intensity-time trace from (m) (black). Hidden Markov Model (HMM) fit used to identify pauses in degradation is shown in red. **o,** Fraction of decay events with detectable pauses. Control traces were obtained from non-degraded SERPs in cells treated with MZB (see Methods) (n=113 traces from 4 independent repeats). **(j, o),** Unpaired Student’s t-test was used for statistical analysis. **** denotes p < 0.0001.

### *In vivo* proteasome degradation kinetics

Human U2OS cells expressing the socRNAs described above were followed over time by spinning disk confocal microscopy. Tracking of single SERPs revealed frequent events in which single GFP foci suddenly decreased in fluorescence intensity, typically over a period of several minutes (Fig. 1d,e; Supplementary Video 1). Treatment with the proteasome inhibitors MG132 or Marizomib (MZB) abolished these decay events (Fig. 1f and Supplementary Fig. 1d), demonstrating that the observed gradual loss in fluorescence reflects proteasomal degradation. To calculate absolute decay rates (in amino acids per second) from intensity-time traces of single SERP decay events, we normalized fluorescence intensities of SERPs using reference fluorescence foci containing a defined number of GFP molecules (See Methods). Using this calibration, an average proteasomal degradation rate of 16.4 ± 5.4 (mean ± s.d.) amino acids per second (aa/s) was calculated (Fig. 1g), which is in agreement with previous measurements of proteasome translocation rates from *in vitro* assays (∼10–30 aa/s)^9,27,28^. Degradation rates were constant over time and did not depend on SERP length (Fig. 1e and Supplementary Fig. 1e), indicating that the very long nature of SERPs does not affect decay kinetics. In our assay, SERPs are fluorescently labeled by binding of the SunTag antibody (STAb)-mStayGold to SunTag peptides (Fig. 1b). To test whether STAb-mStayGold binding to the SERP affects the degradation rate, we reduced labeling density of the SunTag peptide epitopes through reduced STAb-mStayGold expression. Degradation rates were not detectably affected by reduced SunTag labeling stoichiometry (Fig. 1h and Supplementary Fig. 1f,g), indicating that binding of the STAb to the substrate has minimal impact on proteasome decay rates of the SERPs. These results suggest that the proteasome efficiently removes substrate-interacting proteins during degradation *in vivo*. To further confirm this conclusion, degradation rates were determined for SERPs consisting of a string of mNeonGreen molecules (a bright green fluorescent protein) that are intrinsically fluorescent and are not bound by any interacting proteins. These mNeonGreen-SERPs were tethered to the plasma membrane through attachment of a 5×ALFA tag array to the N-terminus (but not the repeat unit) of the SERP (Fig. 1i, Supplementary Fig. 1h and see Methods). Degradation of mNeonGreen-SERPs was induced using the let-7a system, as before. Using this system, an average degradation rate of ∼14.4 ± 7.4 aa/s (mean ± s.d.) was observed (Fig. 1j), in good agreement with the SunTag-based measurements. Together, these results show that STAb-GFP binding to the SERP does not impact decay rate measurements and that our assay therefore enables reliable, quantitative measurements of protein degradation rates by single proteasomes in living cells. As the SunTag labeling system provided improved fluorescent signal and thus more accurate measurements (Supplementary Fig. 1i,j), we continued with the SunTag-based SERPs. Notably, these experiments also reveal that the proteasome can rapidly and efficiently remove even high affinity (picomolar^29^) substrate-interacting proteins during substrate degradation *in vivo*, which contrasts *in vitro* studies where high affinity substrate interactors inhibited substrate decay^30^, highlighting the importance of performing high resolution measurements in cells.

We next assessed processivity of the proteasome (i.e. the number of amino acids that is degraded before the degradation is aborted), a parameter of proteasome activity that is especially difficult to interrogate in live cells using existing methods. We found that most degradation events continued until SERP decay was completed, indicating that the entire SERP polypeptide can be degraded in a single substrate engagement event by the proteasome. However, in a subset of events, SERP fluorescence intensity initially decreased, but this decrease was abruptly halted and followed by the onset of a plateau in the fluorescence intensity (Fig. 1k and Supplementary Video 2), indicative of proteasome stalling or substrate release before degradation was completed. From all degradation events, ∼30% of decay events had aborted after degradation of 10,000 amino acids, resulting in a calculated processivity of ∼15,000 amino acids (Fig. 1l) (i.e. the number of amino acids degraded at which 50% of decay events had aborted). Considering the average protein length of 375 amino acids in the human genome, the measurements show that ∼2–3% of protein decay events are expected to result in incomplete protein degradation, with higher percentages for longer proteins. Such incomplete degradation events may be problematic, as the ubiquitin chain is likely removed during the first substrate engagement of the proteasome, making re-engagement of the proteasome with the degradation intermediate less favorable. Thus, once engaged with their substrates, proteasomes are highly processive machines *in vivo*, which is likely important to limit formation of stable and potentially toxic degradation intermediates.

Visual inspection of intensity traces suggested that some degradation events were interrupted by pauses lasting several minutes (Fig. 1m). Such pauses likely reflect proteasomes that are temporarily stalled during substrate processing. To quantify such proteasome pausing, we applied Hidden Markov Modeling (HMM) to fluorescence intensity-time traces (Fig. 1n). To control for technical noise in intensity-time traces, which could result in false positive pause detection, we generated a negative control dataset with similar technical noise from plateau intensity traces obtained from cells treated with MZB (See Methods). Using this approach, pauses were detected in ∼5% of all degradation traces, corresponding to a pause frequency of around 1/166,000 degraded amino acids (Fig. 1o). This value provides a lower limit of pause frequency, as additional shorter pauses may occur that fall below the detection threshold of our analysis (several minutes). We conclude that proteasomal degradation *in vivo* is fast and highly processive, displaying infrequent, long-lived pauses.

### Impact of substrate diversity on degradation kinetics

*In vitro* work has revealed that substrate fold can profoundly impact proteasome degradation^27,31,32^. We therefore examined how substrate fold and sequence influence proteasome decay kinetics *in vivo*. To compare decay of folded and unfolded protein domains, we engineered SERPs containing either a long unstructured segment derived from cytochrome b ^33^ or the folded SNAP-tag within the repeating unit (Fig. 2a). Somewhat surprisingly, degradation rates were indistinguishable between the folded and unfolded substrates (Fig. 2b), indicating that protein unfolding does not detectably slow substrate translocation on the proteasome *in vivo* under these conditions.

**Fig. 2.**
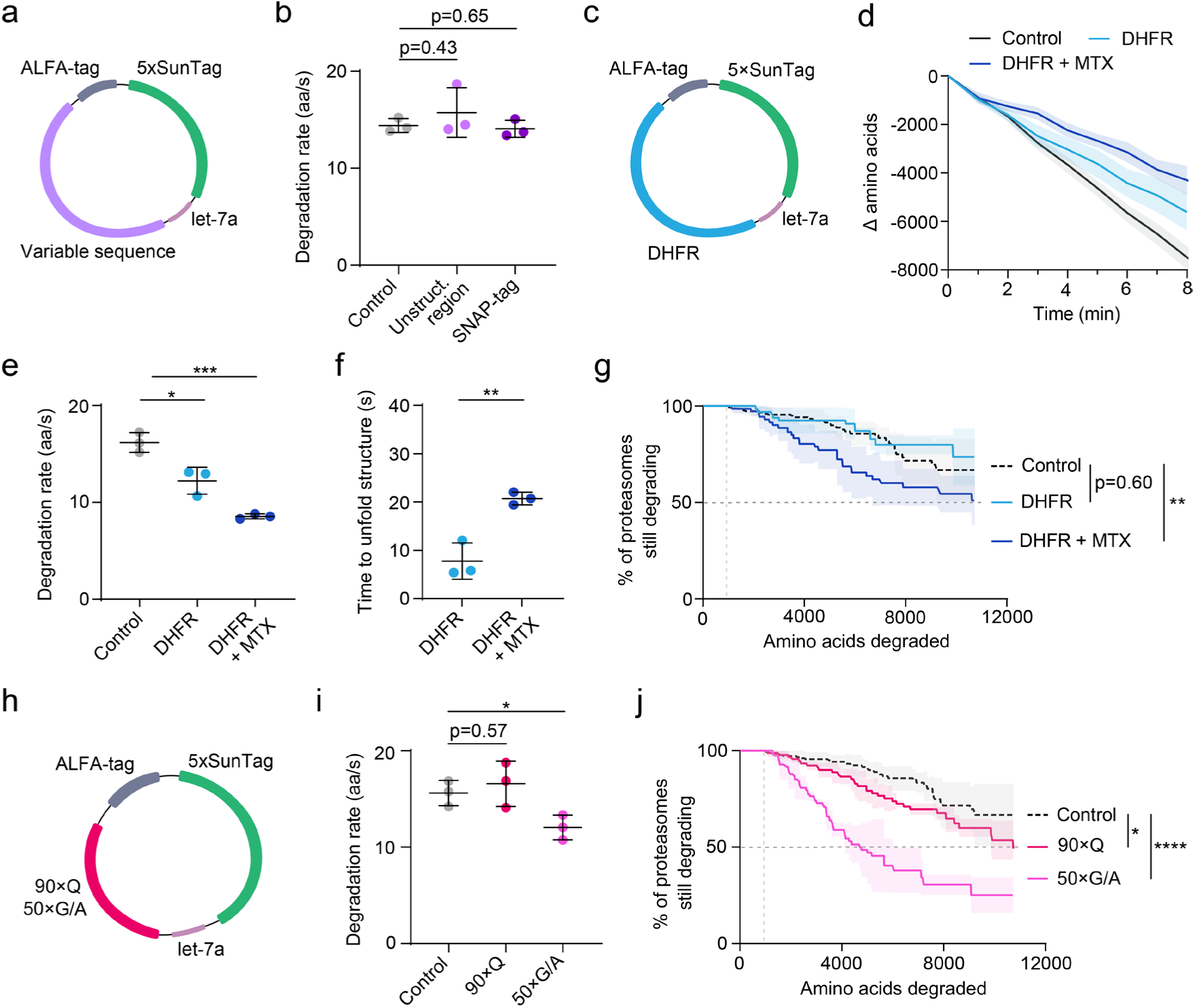
Impact of substrate diversity on degradation kinetics. **a,** Schematic of socRNAs encoding variable test sequence (size of variable sequences are 157 aa (unstructured) and 187 aa (SNAP-tag). Remainder of the sequence is 185 aa). **b,** Comparison of proteasome degradation rate for different SERPs produced from socRNAs depicted in (a) and Fig. 1c (Control). Each data point represents the median from an independent experiment. Horizontal line and error bars represent mean ± s.d of medians (n = 44 traces (control), n = 46 traces (unstructured region) and n = 44 traces (SNAP-tag) from 3 independent repeats). **c,** Schematic of socRNA encoding DHFR (DHFR is 186 aa, the remainder of the sequence is 185 aa). **d,** Average number of amino acids degraded over time for different SERPs produced from socRNAs depicted in (c) (DHFR) and Fig. 1c (Control) with or without MTX treatment. Lines and shaded regions indicate mean ± s.d (n = 11 traces (control), n = 10 traces (DHFR) and n = 9 traces (DHFR + MTX) from 2-3 independent repeats). **e,** Comparison of degradation rate of indicated SERPs and treatment condition. Each data point represents the median from an independent experiment. Horizontal line and error bars represent mean ± s.d of medians (n = 24 traces (control), n = 61 traces (DHFR) and n = 61 traces (DHFR + MTX) from 3 independent repeats). **f,** Time required for unfolding of a single DHFR domain by the proteasome with or without MTX treatment. Each data point represents the median from an independent experiment. Horizontal line and error bars represent mean ± s.d of medians (n = 61 traces (DHFR) and n = 61 traces (DHFR + MTX) from 3 independent repeats). **g**, Kaplan-Meier survival curves representing proteasome processivity on indicated SERPs. Line and shaded region indicate mean ± s.d (n = 170 traces (control), n = 61 traces (DHFR) and n = 61 traces (DHFR + MTX) from 3-5 independent repeats). **h,** Schematic of socRNA encoding homopolymeric amino acid repeats (size of sequences is 90 aa (90×Q) and 50 aa (50×G/A). Remainder of the sequence is 187 aa). **i,** Comparison of proteasome degradation rate for different SERPs produced from socRNAs depicted in (h) and Fig. 1c (Control). Each data point represents the median from an independent experiment. Horizontal line and error bars represent mean ± s.d of medians (n = 122 traces (control), n = 168 traces (90×Q) and n = 136 traces (50×G/A) from 3 independent repeats). **j**, Kaplan-Meier survival curves representing proteasome processivity on indicated SERPs. Line and shaded region indicate mean ± s.d (n = 170 traces (control), n = 168 traces (90×Q) and n = 136 traces (50×G/A) from 3-5 independent repeats). **(b, e, f, i),** Unpaired Student’s t-test was used for statistical analysis. *, **, ***, and **** denote p < 0.05, 0.01, 0.001, and 0.0001, respectively. **(g, j),** Log-rank (Mantel-Cox) test was used for statistical analysis. *, ** and **** denote p< 0.05, 0.01 and 0.0001, respectively.

To explore the limits of protein folding stability on substrate translocation, an extremely tightly folded domain, murine dihydrofolate reductase (DHFR), was inserted into the SERP (Fig. 2c). DHFR structural stability can be further increased by addition of the small molecule methotrexate (MTX) to create a substrate that strongly inhibits the proteasome *in vitro*^30^. While DHFR induced a brief pause during degradation (7 s) that was prolonged upon MTX addition (20 s), the proteasome nevertheless degraded this highly stable domain with surprising efficiency (Fig. 2d–f). Even more surprisingly, processivity of decay was largely unaffected for decay of DHFR domains, with only a slight decrease upon MTX treatment (Fig. 2g). From these processivity measurements we could calculate that only in 1.7% of cases, the proteasome aborted degradation when encountering a MTX-stabilized DHFR domain. These data indicate that *in vivo* the proteasome can unfold even highly stable protein domains, and can remain engaged with the substrate even when paused to unfold such protein domains.

We next focused on other notoriously difficult proteasome substrates, homopolymeric poly-glutamine (polyQ) and glycine/alanine tracts. Long polyQ tracts occur by genetic CAG repeat expansion and underlie several neurodegenerative disorders, including Huntington’s disease, with polyQ lengths of ∼36 residues or greater typically being associated with disease^34^. PolyQ repeat containing proteins in neurodegenerative disease often persist in disease, despite being ubiquitinated and associated with the proteasome^35,36^. Consistently, several in *vitro* studies using purified components have reported that long polyQ tracts can resist proteasomal degradation^37-39^, although others have found efficient degradation *in vitro*^40^. To measure decay rates of polyQ tracts *in vivo*, a polyQ tract of 90 residues was inserted into the SERP (Fig. 2h), a polyQ length that is associated with severe, juvenile Huntington disease onset^34^. Surprisingly, even these very long polyQ tracts had no detectable effect on average degradation speed and only a very small effect on processivity (Fig. 2i,j), indicating that long polyQ stretches do not inherently impede proteasome degradation. In contrast, a 50× glycine/alanine (G/A) repeat derived from Epstein–Barr virus, another sequence that is known to impede proteasomal degradation^41^, caused a significant reduction in both speed and processivity (Fig. 2i, j), confirming the sensitivity of the SERP assay. These results show that long polyQ segments can be efficiently degraded *in vivo*, and suggest that the inability to degrade polyQ proteins in disease may be due to other protein or cell features.

### Directionality of proteasomal degradation

In the assay described above, protein degradation is triggered by C-terminal ubiquitination of SERPs through a ribosome quality control pathway, resulting in degradation in the C-to-N direction. Proteasomes, however, are also known to engage protein substrates from the N-terminus or at substrate-internal sites^8,42^, but it is unclear whether degradation directionality affects proteasome degradation kinetics. To study N-to-C proteasomal degradation, a single copy of a N-terminal poly-leucine sequence was introduced into the SERP (see Supplementary Fig. 2a,b and Methods), which acted as a potent degron and led to efficient SERP degradation (Fig. 3a,b and Supplementary Fig. 2a-c). Degradation of SERPs with the N-terminal degron was abolished by proteasome inhibition (Fig. 3c and Supplementary Fig. 3c). Interestingly, in the subset of cases where SERP degradation was incomplete, fluorescence intensity of SERPs increased again after degradation was aborted, indicating that SERP degradation occurred mostly co-translationally and confirming that degradation occurred in N-to-C (as the C-terminus of the SERP is bound to the tRNA inside the translating ribosome) (Supplementary Fig. 2d). Such increase in SERP foci intensity upon incomplete degradation was rarely observed in case of the C-to-N decay assay (Supplementary Fig. 2e,f), validating the directionality of both decay assays.

**Fig. 3.**
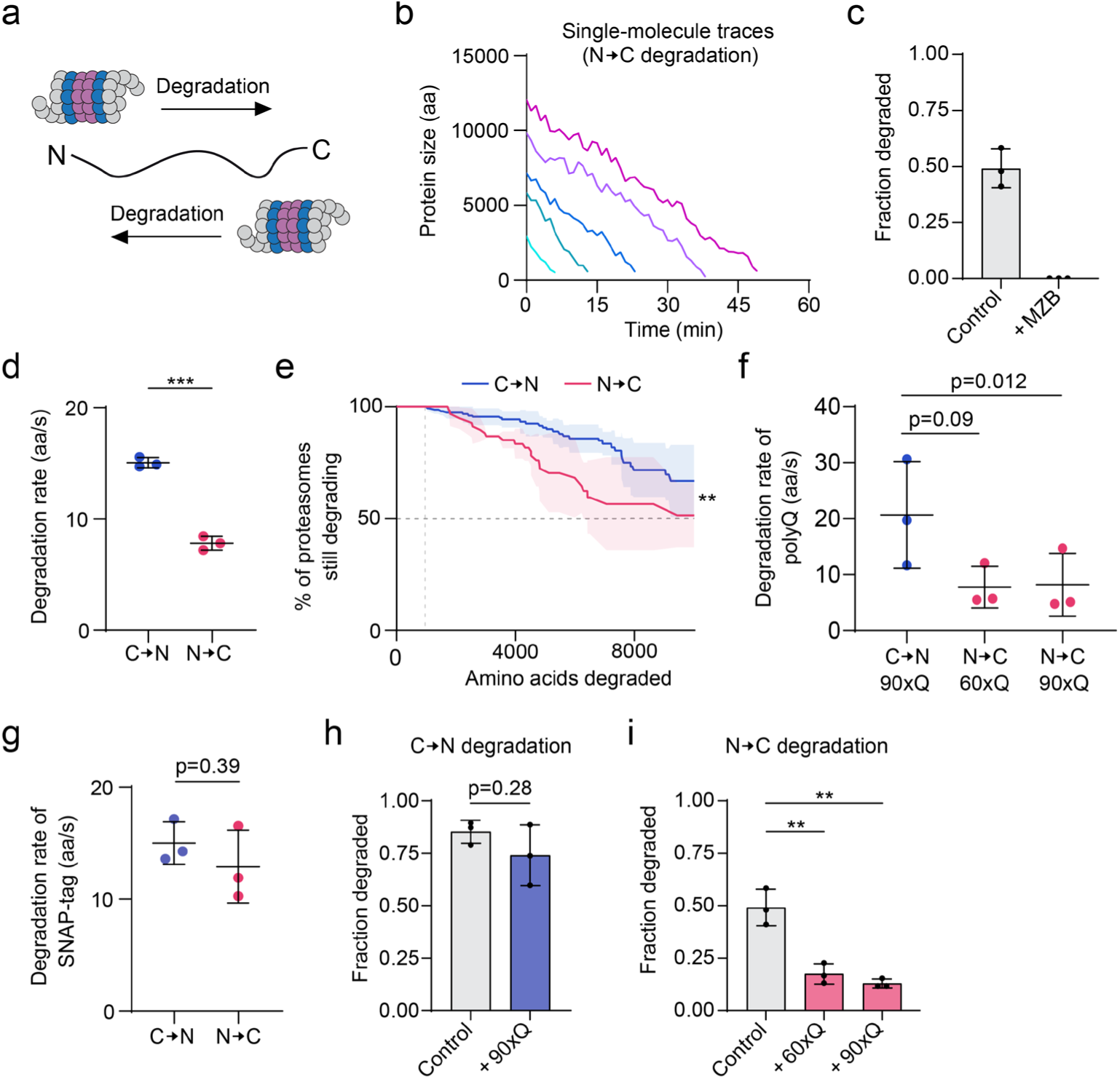
Effect of proteasome directionality on degradation. **a**, Schematic depicting proteasomal degradation from N-to-C terminus and C-to-N terminus. **b,** Representative fluorescence intensity-time traces of N-to-C degradation events. **c,** Fractions of SERPs subject to degradation in control vs. MZB-treated condition (n = 130 traces (Control) and n = 37 traces (+MZB) from 3 independent repeats). **d,** Proteasomal degradation rates for SERPs degraded with either C-to-N and N-to-C directionality. SERP sequence was identical in both conditions. SERPs degraded from C-to-N or N-to-C were expressed from socRNAs shown in Fig. 1c and Supplementary Fig. 2a, respectively. Each data point represents the median from an independent experiment. Horizontal line and error bars represent mean ± s.d of medians (n = 79 traces (C-to-N) and 62 traces (N-to-C) from 3 independent repeats). **e**, Kaplan-Meier survival curves representing proteasome processivity for C-to-N and N-to-C degradation. SERP sequence was identical in both conditions. Lines and shaded region indicate mean ± s.d. Blue line (C-to-N) was replotted from Fig. 1l for comparison. Log-rank (Mantel-Cox) test was used for statistical analysis. ** denotes p<0.01 (n = 170 traces (C-to-N) and 62 traces (N-to-C) from 3 independent repeats). **f,** Comparison of proteasomal degradation rate of the polyQ sequence for C-to-N and N-to-C degradation. Each data point represents the median from an independent experiment. Horizontal line and error bars represent mean ± s.d of medians (n = 168 traces (C-to-N, 90×Q), n = 15 traces (N-to-C, 60xQ) and 13 traces (N-to-C, 90×Q) from 3 independent repeats). **g,** Comparison of proteasomal degradation rate from C-to-N and N-to-C on SNAP-tag sequence alone (see Methods). Each data point represents the median from an independent experiment. Horizontal line and error bars represent mean ± s.d of medians (n = 44 traces (C-to-N) and 71 traces (N-to-C) from 3 independent repeats). **h,** Fraction of SERPs undergoing degradation in C-to-N decay assay (n = 136 traces (Control) and 138 traces (+90×Q) from 3 independent repeats). **i,** Fraction of SERPs undergoing degradation in N-to-C decay assay. Control data were replotted from panel (c) (n = 130 traces (Control), 111 traces (+60×Q) and 58 traces (+90×Q) from 3 independent repeats). **(d, f, g, h, i,)** Unpaired Student’s t-test was used for statistical analysis. ** and *** denote p < 0.01 and 0.001, respectively.

Comparison of degradation rates for C-to-N and N-to-C degradation for SERPs with identical amino acid sequences revealed a surprising difference; proteasomal degradation from the C-to-N proceeded at ∼15 aa/s, whereas degradation from the N-to-C proceeded at only ∼8 aa/s (Fig. 3d). Proteasome processivity was also reduced when degradation occurred in the N-to-C direction (Fig. 3e). To assess whether the impact of the directionality on substrate processing represents a general feature underlying proteasomal degradation, we examined two additional substrate types. We introduced either the SNAP-tag or the polyQ tract into the SERP repeat unit and compared degradation rates for both decay directionalities. Intriguingly, the directional asymmetry was different for different substrates, with polyQ sequences also showing a strongly reduced decay rate for N-to-C compared to C-to-N decay directionality (Fig. 3f). In contrast, only a very modest reduction in decay rate (which did not reach statistical significance) was observed for the SNAP-tag SERP (Fig. 3g). Surprisingly, for polyQ-containing SERPs we also observed differences in the *fraction* of substrates for which degradation was (detectably) initiated, an effect that was not observed for C-to-N degradation of polyQ substrates (Fig. 3h,i). An unrelated unstructured sequence of the same length did not impact the fraction of substrates that was degraded in the N-to-C assay (Supplementary Fig. 2g). These observations reveal that proteasomal degradation kinetics are substantially impacted by the direction of degradation, and thus by the mode of substrate engagement, in a substrate specific manner. Moreover, these data illustrate the importance of measuring different substrate processing parameters, e.g. proteasome recruitment, decay rate, processivity and directionality, independently to disentangle how sequences like polyQ tracts impact protein degradation efficiency.

### Proteasome-ribosome collisions

In addition to differences in the site of substrate engagement, previous studies have also suggested that proteasomes can engage with protein substrates at different times in the protein’s life-cycle, including co-translationally in a process termed co-translational protein decay (CTPD)^43^. Consistent with this, nascent chains are known to be ubiquitinated co-translationally^44^, and proteasomes have been reported to associate with translating ribosomes^44-46^. In this context it is interesting to note that our results show that proteasomes degrade proteins at a substantially higher rate than ribosomes synthesize them (∼8-15 vs. 2-3 aa/s ^22,47^), suggesting that proteasomes can catch up and collide with translating ribosomes. Using protein synthesis and decay rates we calculated that such collisions are expected to occur if the proteasome engages with the nascent polypeptide while the ribosome is translating the first ∼67% of the coding sequence (See Methods). Previous work on ribosome-ribosome collisions has identified specialized processes that deal with such collisions^48^, but the consequences of proteasome-ribosome collisions are unknown.

We considered three possible outcomes of proteasome-ribosome collisions (Fig. 4a); first, the proteasome releases the substrate upon collision and the ribosome continues translating the mRNA. Second, the proteasome trails the translating ribosome, continuously degrading the nascent chain as it emerges from the ribosome. In the third model, proteasome-ribosome collisions trigger dissociation of the ribosome from the mRNA, as is the case for ribosome-ribosome collisions^48^. We first tested if proteasomes release their substrate upon collision with a ribosome. For this analysis, we focused on co-translational decay events on socRNAs translated by more than one ribosome, as in these cases the socRNA can still be tracked upon complete degradation of one of the nascent polypeptides as the GFP spot does not completely disappear due to the presence of the second nascent chain within the same spot (Fig. 4b). We reasoned that if a proteasome dissociates upon collision with a ribosome (Model 1), the rate of GFP spot intensity increase *after the decay event* should be identical to the intensity increase *before the decay event*, as the number of ribosomes on the socRNA that contribute to GFP signal production remains constant (Fig. 4c and Supplementary Fig. 3a,b). In contrast, if proteasomes continuously trail ribosomes (Model 2) or induce ribosome dissociation (Model 3), the GFP intensity slope after the decay event should be reduced compared to the initial rate, as in both Models 2 and 3 the ribosome associated with the decay event no longer contributes to the GFP intensity increase (Fig. 4c and Supplementary Fig. 3a). Importantly, in the socRNA assay, no new ribosomes are recruited to translating socRNAs^22^, simplifying interpretation of any increase in the rate of GFP accumulation. In 29 out of 36 (∼80.6%) events, the rate of GFP intensity increase was reduced after the decay event (Fig. 4d,e), suggesting proteasome release is not a common outcome of proteasome-ribosome collisions.

**Fig. 4.**
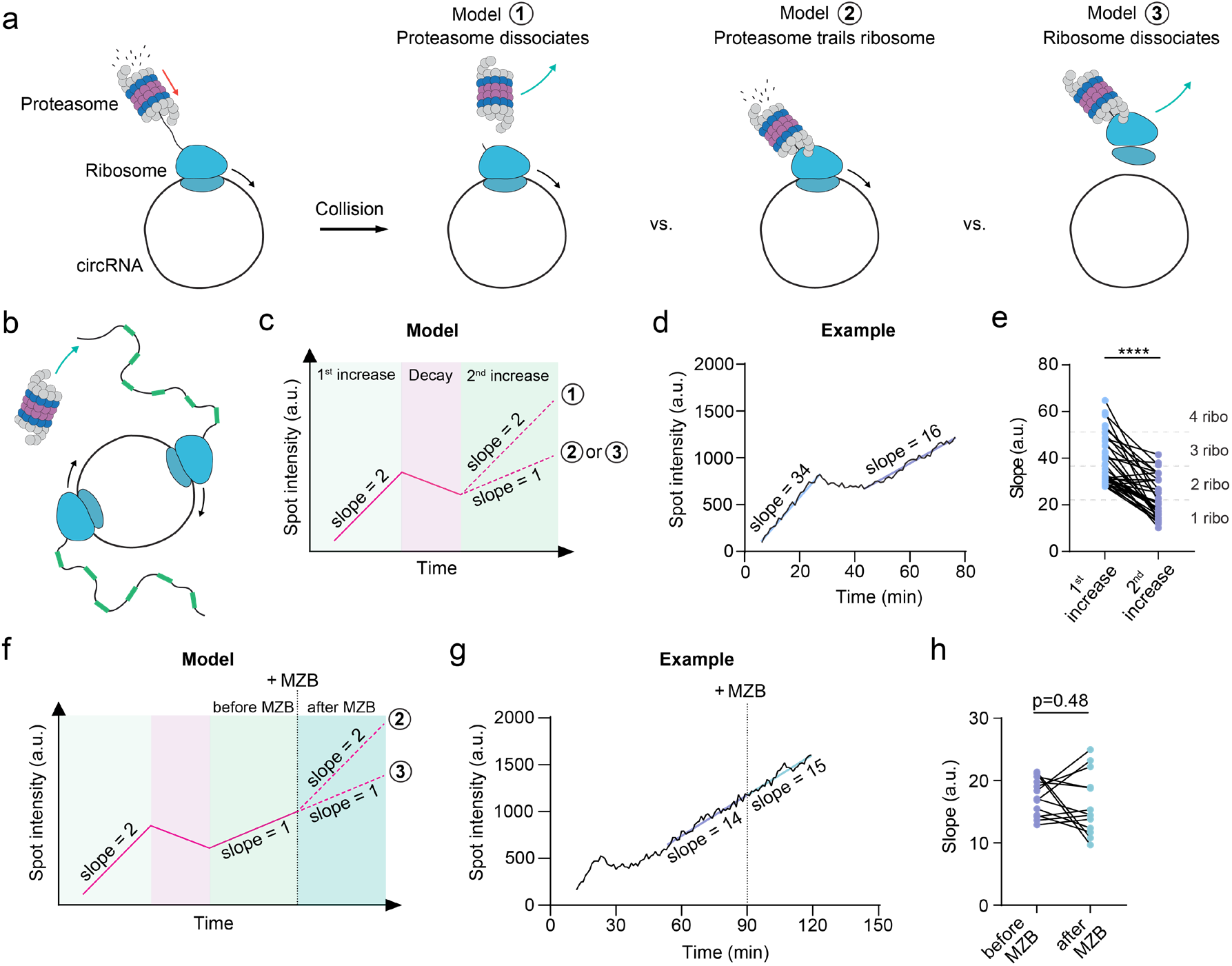
Proteasome-ribosome collisions promote ribosome dissociation. **a**, Models of different outcomes of proteasome-ribosome collisions. Upon collision, the proteasome either dissociates (Model 1), trails the translating ribosome while continuously degrading the newly synthesized polypeptide (Model 2), or the ribosome dissociates from the mRNA (Model 3). **b,** Schematic of two ribosomes translating a single socRNA. A proteasome engages one of the two nascent chains. Prior to degradation, both nascent chains exhibit identical fluorescence intensities. **c,** Schematic of SERP foci intensity over time in the case of the different models described in (a). Note that in all models, socRNAs are translated by two ribosomes. During the first 1^st^ increase phase (synthesis of SERPs by ribosomes), the slope reflects translation by two ribosomes on the same socRNA. In the decay phase, one nascent chain is being co-translationally degraded, while the other continues to be synthesized. Since degradation occurs at a higher rate, the total SERP intensity decreases. The second increase phase either shows a similar slope as the original slope (Model 1, proteasome dissociation) or shows a reduced slope (Models 2 or 3, proteasome trailing the ribosome or ribosome dissociation). **d,** Representative intensity-time trace showing initial translation by two ribosomes, followed by signal decay corresponding to co-translational degradation of one nascent chain, and a subsequent increase in signal, which occurs at a lower rate (reduced slope), reflecting continued SERP synthesis by fewer ribosomes. **e,** Comparison of the slopes of intensity-time traces before and after the decay event. Only traces were included in which the 1^st^ increase phase was consistent with translation by ≥2 ribosomes. Dashed lines reflect the expected number of ribosomes based on fluorescence intensities (n = 36 traces from 3 independent repeats). **f,** Schematic of the strategy used to discriminate between Model 2 (proteasome trails ribosome) and Model 3 (ribosome dissociates upon collision). The proteasome inhibitor MZB is added during the second increase phase to discriminate the two models based on slopes of the second increase phase. **g,** Representative intensity-time trace showing co-translational degradation of one nascent chain on a socRNA translated by two ribosomes. MZB was added during the 2^nd^ increase phase. Note that the slope is not altered upon MZB treatment. **h,** Comparison of the slopes of the 2^nd^ increase phase before and after MZB addition. While some changes are observed, likely due to technical noise caused by the relatively short intensity-time traces in this experimental setup, the average slope is unaffected (n = 13 traces from 3 independent repeats). **(e, h)** Paired Student’s t-test was used for statistical analysis. **** denotes p < 0.0001.

To discriminate between continuous trailing of the ribosome by the proteasome (Model 2) and ribosome dissociation (Model 3), we asked if acute proteasome inhibition by MZB after a decay event is completed would increase GFP accumulation rates, as is predicted in the proteasome trailing model (Fig. 4f and Supplementary Fig. 3c). As a control, addition of MZB to *ongoing* degradation events (i.e. those that had not yet completed, so no proteasome-ribosome collision had yet occurred) led to a rapid increase in GFP accumulation rates, indicating that MZB treatment efficiently inhibits proteasomes actively degrading in a N-to-C direction as well (Supplementary Fig. 3d, see Methods). Comparison of GFP slopes before and after acute proteasome inhibition revealed no significant increase in the slope (Fig. 4g,h and Supplementary Fig. 3d), suggesting that the ribosome associated with the co-translational decay event had ceased translation and dissociated from the mRNA. These results support a model in which proteasome–ribosome collisions during CTPD induce ribosome removal from the mRNA, possibly reflecting a novel co-translational quality control pathway.

### Role of p97 in proteasomal decay

Proteasomes are thought to function together with a network of regulatory factors important for substrate degradation. A key regulatory factor is p97/VCP (hereafter referred to as p97), an AAA+ family ATPase that can extract and unfold ubiquitinated substrates from macromolecular complexes, membranes or chromatin^15^. p97 has been intimately linked to proteasome-dependent protein degradation, but which substrates p97 acts on and how it promotes proteasomal degradation is still debated^49,50^. To probe which aspects of proteasomal function are controlled by p97, we inhibited p97 using the small molecule inhibitor CB5083^51^ and assessed substrate processing kinetics. We first examined N-to-C degradation dynamics in the absence of p97 activity, but found that p97 inhibition did not substantially alter either the fraction of degraded SERPs, the rate of decay or proteasome processivity (Supplementary Fig. 4a-c). Similarly, for the C-to-N degradation assay, in which degradation is triggered by ribosome quality control, the rate and processivity of degradation was not affected by p97 inhibition (Fig. 5a,b). However, initiation of protein degradation was substantially delayed upon p97 inhibition in this assay, as seen by a long plateau phase in the GFP intensity between the end of the translation phase and the start of the decay phase, consistent with the known role for p97 in ribosome quality control in extracting ubiquitinated polypeptides from 60S ribosomes to allow their degradation^52^ (Fig. 5d-e and Supplementary Fig. 4d). Somewhat surprisingly, in the majority of cases, degradation did eventually ensue even when p97 was inhibited, possibly due to incomplete inhibition of p97, or through a redundant pathway in nascent chain extraction from the ribosome. Together, these results uncover the kinetics of ribosome quality control, and importantly, show that p97 activity is not generally required for proteasomal degradation.

**Fig. 5.**
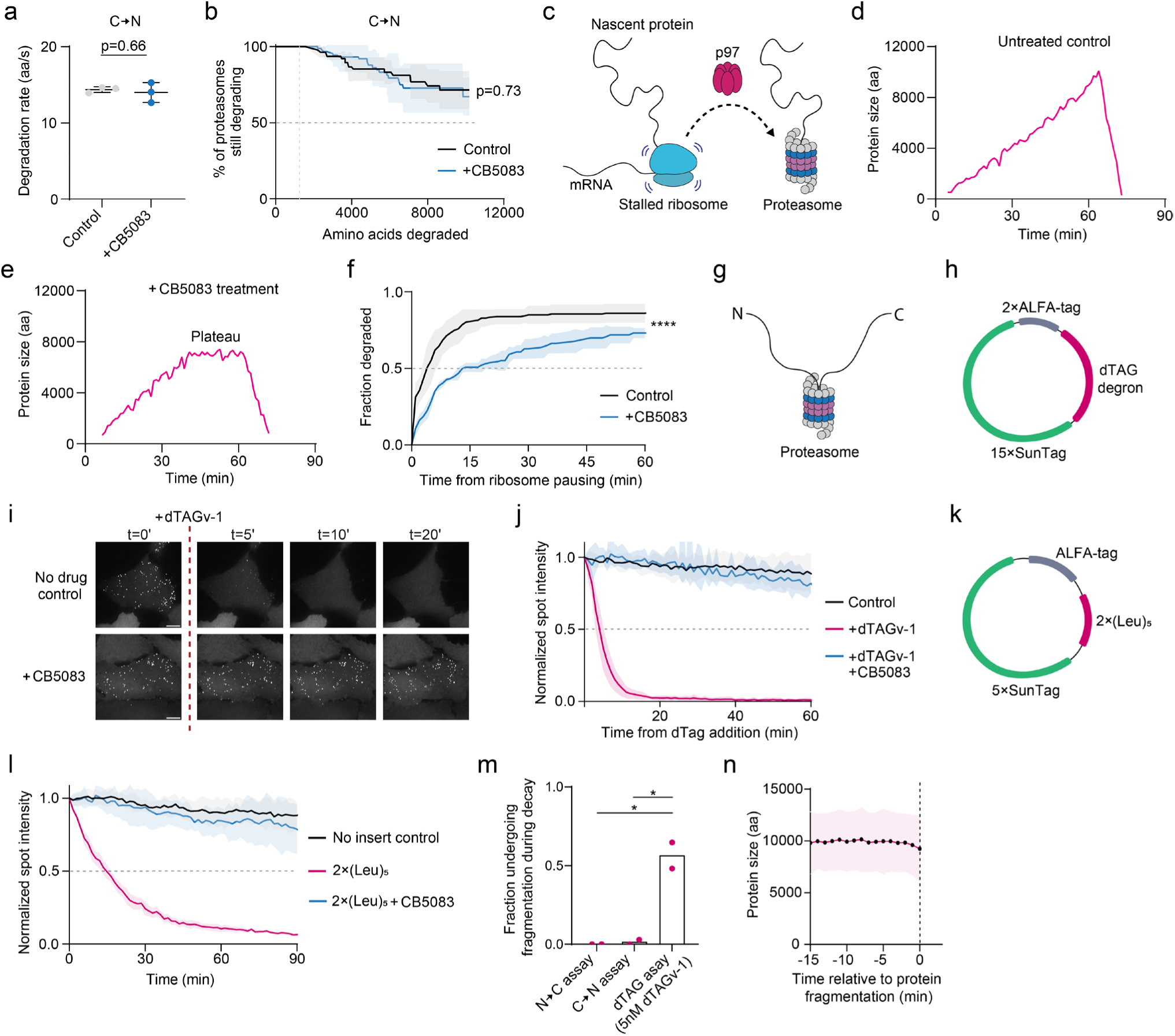
p97 is required for substrate-internal protein degradation. **a,** Analysis of proteasomal degradation rate for C-to-N directionality in the presence or absence of the p97 inhibitor CB5083. Each data point represents the median from an independent experiment. Horizontal line and error bars represent mean ± s.d of medians (n = 63 traces (control) and 95 traces (+CB5083) from 3 independent repeats). **b,** Kaplan-Meier survival curves representing proteasome processivity for C-to-N degradation in the presence or absence of CB5083. Line and shaded region indicate mean ± s.d (n = 63 traces (control) and 95 traces (+CB5083) from 3 independent repeats). **c,** Schematic illustrating the role of p97 in extracting the nascent polypeptide from the ribosome to allow proteasome engagement with the nascent polypeptide. **d,e,** Representative intensity-time trace for a SERP undergoing synthesis and subsequent decay upon ribosome quality control. The plateau phase represents the time from ribosome stalling at the end of the cleaved mRNA to degradation by the proteasome. Untreated cell (d) and cell treated with the p97 inhibitor CB5083 (e) are shown. The pronounced plateau phase upon p97 inhibition is likely caused by slow extraction of the nascent chain from the 60S ribosome. **f,** Cumulative incidence curve showing the fraction of SERPs that has initiated degradation relative to the moment of ribosome stalling (i.e. plateau onset) (n = 167 traces (control) and 95 traces (+CB5083) from 3-6 independent repeats). **g,** Schematic illustrating internal substrate engagement by the proteasome. **h,** Schematic of the socRNA used for internal substrate recruitment experiments. The socRNA encodes 15×SunTag repeats, 2×ALFA-tag repeats, and a single dTAG-inducible degron. **i,** Cells expressing the internal dTAG SERP shown in (h) were treated with 500 nM dTAG ligand (dTAGv-1) in the presence or absence of the p97 inhibitor CB5083. Scale bars, 10 μm. **j,** Quantification of SERP degradation kinetics following addition of 500 nM dTAGv-1. Pre-treatment of cells with p97 inhibitor CB5083 prevents degradation. Line and shaded region indicate mean ± s.d (n = 10 fields of view (control), n = 8 fields of view (+dTAGv-1) and n = 7 fields of view (+dTAGv-1+CB5083) from 2-3 independent repeats). **k,** Design of socRNA used to assess internal degradation via poly-leucine degrons, which is made up of two stretches of five consecutive leucine residues. **l,** Quantification of degradation kinetics of polyleucine-containing SERPs encoded by the socRNA construct shown in (k). Pre-treatment of cells with VCP inhibitor CB5083 prevents degradation. Line and shaded region indicate mean ± s.d (n = 10 fields of view (control), n = 8 fields of view (2x(Leu)_5_) and n = 8 fields of view (2x(Leu)_5_+CB5083) from 2-3 independent repeats). **m,** Fraction of SERPs undergoing fragmentation during the degradation phase across indicated assays. Each data point represents the median from an independent experiment (n = 41 traces (N-to-C assay), n = 55 traces (C-to-N assay) and n = 44 traces (5 nM dTAGv-1) from 2 independent repeats). **n,** Average SERP size aligned to the moment of protein fragmentation in cells treated with 5 nM dTAGv-1. Dashed line represents the moment of SERP fragmentation. SERPs were produced as shown in (h). Only events that exhibited fragmentation were included in the analysis. Line and shaded region indicate mean ± s.d (n = 23 traces from 2 independent repeats). **(a, m),** Unpaired Student’s t-test was used for statistical analysis. * denotes p < 0.05. **(b, f),** Log-rank (Mantel-Cox) test was used for statistical analysis. **** denotes p<0.0001.

In addition to N- or C-terminal engagement, proteasomes can engage substrates internally as well^14,33^ (Fig. 5g). However, how proteasomes are recruited internally and whether substrate processing occurs with similar kinetics for internal engagement is not well understood. To induce substrate-internal engagement by the proteasome, we engineered internal ubiquitination sites all along the SERP. To this end, the inducible dTAG degron, which recruits a potent E3 ubiquitin ligase upon ligand addition, was inserted into the repeat unit of the SERP (Fig. 5h). Addition of the dTAG ligand (dTAGv-1) induced rapid SERP degradation, with most GFP foci disappearing within minutes (Fig. 5i). Strikingly, inhibition of p97 completely blocked substrate-internal degradation (Fig. 5i,j and Supplementary Video 3), identifying a specific role of p97 in substrate-internal degradation. It is thought that proteasomes require an unstructured sequence to initiate decay^8,33^, and the absence of such unstructured sequence in internal engagement could explain the requirement for p97 in substrate-internal decay. However, even after introduction of long (up to 180 aa) unstructured sequences into the SERP near the dTAG degron, inhibition of p97 strongly inhibited protein decay, indicating that unstructured regions alone are insufficient to bypass the need for p97 in substrate-internal proteasomal decay (Supplementary Fig. 5a,b). To further validate the role of p97 in substrate-internal protein decay, we replaced the dTAG degron with an internal poly-leucine degron (Fig. 5k), which also acted as a potent destabilizing sequence (see Supplementary Fig. 2a-c). SERPs containing the internal poly-leucine degron were rapidly degraded in untreated cells, but were completely resistant to decay upon p97 inhibition (Fig. 5l). Together, these results show that substrate processing depends heavily on the mode of substrate engagement, and identify a selective role for p97 in substrate-internal engagement by the proteasome.

To better understand the role of p97 in substrate-internal engagement of proteasomes, we examined protein degradation kinetics for substrate-internal engagement in more detail. Previous structural and biochemical work has suggested that the central channel of the proteasome can accommodate two polypeptide chains simultaneously^53,54^, but it is unclear if both chains are processively and efficiently processed *in vivo*. To assess internal engagement by single proteasomes, we first optimized the internal engagement assay. In the dTAG assay, the SERP may be ubiquitinated at multiple sites along the polypeptide, so it is possible that multiple proteasomes engage single SERPs simultaneously, making interpretation of decay kinetics more challenging. To ensure recruitment of just a single proteasome to each SERP, we reduced dTAGv-1 concentration and found that at 5 nM ligand only a small subset of SERPs was degraded (Supplementary Fig. 5c), greatly reducing the probability that a single SERP is degraded by multiple proteasomes simultaneously. Surprisingly, when analyzing internal engagement events by single proteasomes, we found that many processive degradation events (∼55%) were preceded by substrate fragmentation, in which a single SERP spot split into two spots (Fig. 5m and Supplementary Fig. 5d,e). In most cases, one of the two SERP fragments underwent decay immediately after substrate fragmentation, while the other remained stable (Supplementary Fig. 5d-f), suggesting that the proteasome released one substrate fragment while continuing to degrade the other.

Interestingly, processive substrate degradation of the SERP fragment that was degraded after fragmentation occurred at ∼15 aa/s (Supplementary Fig. 5g), a rate consistent with C-to-N, but not N-to-C directionality of decay (see Fig. 3). These results suggest that the proteasome preferentially remains associated with the C-terminus while letting go of the N-terminus upon substrate fragmentation, consistent with our findings that degradation from the C-terminus is faster and more processive than degradation from the N-terminus for this substrate. To determine whether limited processive degradation occurred while the proteasome was engaged with both chains simultaneously (i.e. before substrate fragmentation), we plotted GFP intensity over time before substrate fragmentation (Fig. 5m) (See methods). This analysis did not reveal a substantial decrease in GFP intensity before fragmentation, indicating that at our detection threshold no processive degradation of two chains simultaneously could be observed. Together, these results suggest that the proteasome can engage and cleave substrates internally, but is poorly processive while associated with both chains simultaneously, rapidly releasing one of the two chains and degrading the other. Substrate-internal engagement was uniquely dependent on p97, further demonstrating that the mode of substrate engagement determines subsequent substrate processing.

## Discussion

Here, we report a high-resolution live-cell assay to visualize substrate engagement and processing by single proteasomes. This assay provides quantitative measurements of different sub-steps in substrate degradation, including substrate recruitment rates, degradation kinetics and proteasome processivity, revealing detailed mechanistic insights into proteasome activity *in vivo*, and providing a broadly-applicable new technology to study proteasome regulation.

Our measurements show that proteasomes degrade substrates at ∼15 amino acids per second *in vivo,* providing the first such measurements in live cells. This rate is likely set by the translocation rate of the internal proteasomal AAA-protein rather than by the peptide hydrolysis rate. Moreover, we find that proteasomes are highly processive, showing a substrate release probability of just 0.00004 per amino acid. Contrasting *in vitro* results, both the decay rate and processivity were not substantially impacted by physiological variations in protein fold stability or by high-affinity protein-protein interactions of the substrate, demonstrating the very strong translocase activity of the proteasome *in vivo*. Similarly, polyQ tracts, which can resist degradation *in vitro*, showed highly efficient degradation *in vivo* (for C-to-N decay). Possibly, additional co-factors such as p97 or ZFAND proteins, or post-translational modifications not present in *in vitro* reactions help the proteasome in rapid and efficient degradation of tightly folded substrates *in vivo^15,55,56^*.

The proteasome can engage substrates at different sites and translocate substrates in both C-to-N and N-to-C direction. Interestingly, decay rates and processivity are not identical for decay in both directions, with C-to-N decay showing a higher rate and processivity for 2 out of 3 substrates tested. Possibly, protein unfolding rates differ in both decay directions. However, it is unlikely that differences in protein unfolding explain all the results, as 1) we found that the presence of tight protein folds does not slow the proteasome down substantially (Fig. 2b) and 2) polyQ tracts also show a clear asymmetric directionality, even though they are likely not folded. Alternatively, it is possible that the proteasome has an intrinsic C-to-N decay preference, for example due to more favorable interactions of the proteasome RPT subunit pore loops that push the substrate through the central channel with (a subset of) substrate amino acids or with the amino acid backbone. If interactions with amino acid side chains are relevant, the precise amino acid composition of a substrate could determine how strong the C-to-N preference is for substrate translocation. Further exploration of sequence space will be interesting to understand how amino acid identity impacts substrate translocation asymmetry.

The AAA+ family protein p97 has been widely implicated in proteasome-dependent protein degradation, but its precise role has remained unclear. We find that p97 is not needed for degradation from substrate termini, and thus is not a general cofactor of the proteasome. However, p97 is essential for substrate processing upon internal engagement of the substrate by the proteasome. Possibly, p97 unfolds protein-internal regions to allow for proteasome engagement, which may not be needed for recruitment to substrate termini, as these may be more readily accessible.

In this study, we have leveraged our single-proteasome imaging assay to understand how substrate engagement mode impacts substrate processing. However, we envision many additional applications of this assay in studying protein turnover. For example, it will be interesting to understand whether distinct proteasome populations, such as nuclear proteasomes, immunoproteasomes, or proteasomes operating in different cell types or under stress conditions, process substrates differently. Moreover, it will be interesting to further dissect the role of other co-factors, ubiquitin chain architecture and post-translational modifications of substrates and proteasomes on decay kinetics. Even rates of substrate ubiquitination and recruitment to the proteasome can be further explored, as illustrated by our kinetic measurements of protein decay during ribosome quality control.

In summary, we have developed a new assay to visualize proteasome activity *in vivo* with molecular resolution, which has revealed how distinct modes of substrate engagement drive substrate processing with diverse kinetics.

## Methods

### Cell culture

U2OS cells used for imaging experiments were engineered to stably express SunTag antibody (STAb)–mStayGold, ALFA-tag nanobody (ALFANb)-CAAX or ALFANb-Halo-CAAX, and TetR. Cells were maintained at 37 °C and 5% CO₂ in Dulbecco’s Modified Eagle Medium (DMEM; 4.5 g/L glucose, Gibco) supplemented with 10% fetal bovine serum (FBS; Sigma-Aldrich) and 1% penicillin–streptomycin (Gibco). All cell lines were routinely screened and verified to be free of mycoplasma contamination.

### Cell line generation

Stable cell lines were generated via lentiviral transduction. Lentivirus was produced in HEK293T cells by co-transfecting cells at ∼40% confluency with the lentiviral construct of interest and the packaging plasmids pSPAX2 and pMD2.G using polyethylenimine (PEI; Polysciences Inc) as transfection reagent. Medium was refreshed 24 hours post-transfection, and viral supernatant was collected 72 hours after transfection. Lentiviral transduction was performed by plating U2OS in 6-well plates and adding viral supernatant supplemented with 10 μg/mL Polybrene (Santa Cruz Biotechnology Inc), followed by spin infection for 100 minutes at 2,000 rpm at 37 °C. Monoclonal cell lines with homogenous transgene expression levels were generated by FACS-sorting of individual cells into 96-well plates at least 10 days after transduction.

### Drug treatment

SERP expression was induced by a brief pulse of doxycycline (1 μg/mL; Sigma-Aldrich) for 5-8 minutes, followed by two washes with imaging medium to remove residual doxycycline. To examine the impact of proteasome and p97/VCP inhibition on SERP degradation, cells were pre-treated for 10 minutes with either MG132 (10 μM; Merck) or CB5083 (5 μM; MedChemExpress) prior to the start of imaging. For acute proteasome inhibition during live-cell imaging, Marizomib (5 μM; Merck) was added directly to cells, as we found that MG132 did not inhibit the proteasome with rapid kinetics. Unless noted otherwise, degradation of dTAG-containing SERPs was triggered by the addition of 500 nM dTAG-v1 (Bio-Techne). To terminate translation and release nascent SERPs from ribosomes, cells were treated with puromycin (0.1 mg/mL; Thermo Fisher Scientific). For experiments involving DHFR-containing SERPs, methotrexate (MTX; 5 μM, Merck) was added 5 minutes prior to the start of imaging.

### Microscopy

#### Microscope

Unless stated otherwise, fluorescence microscopy experiments were performed using a Nikon TI2 inverted microscope with a large field of view X-light V3 spinning disc (Crest Optics) and an ORCA-Quest2 camera and a motorized piezo stage. For experiments shown in Fig. 1i,j and Supplementary Fig. 1h,i, experiments were performed using a Nikon TI1 inverted microscope equipped with a Yokogawa CSU-X1 spinning disc and an iXon Ultra 897 EM-CCD camera (Andor). A 100× oil immersion objective (NA = 1.49) was used for all acquisitions. NIS Elements perfect focus system was employed to maintain focus during time-lapse acquisitions.

#### Live-cell imaging

Cells stably expressing STAb-mStayGold, ALFANb-CAAX or ALFANb-Halo-CAAX, and TetR were seeded in 96-well glass-bottom plates (Matriplates, Brooks Life Science Systems). The following day, cells were transfected with socRNA constructs using Fugene HD (Promega). Prior to imaging, medium was replaced with CO₂-independent Leibovitz’s L-15 imaging medium (Gibco) supplemented with 5% FBS (Sigma-Aldrich) and 1% penicillin–streptomycin (Gibco). socRNA expression and SERP production was induced using doxycycline (1 μg/mL). Unless noted otherwise, socRNA and SERP expression was induced by two short doxycycline pulses (1 μg/mL): the first administered 1 hour prior to imaging, and the second 20 minutes before imaging onset. Each pulse lasted 5 minutes and was followed by two washes with imaging medium to remove residual doxycycline. The two-pulse strategy enabled the identification of cells efficiently expressing SERPs (first pulse), while allowing real-time tracking of SERPs induced by the second pulse. For substrate-internal proteasome degradation experiments (Fig. 5i-n, Supplementary Fig. 4f-k), expression was induced by a single 8-minute doxycycline pulse. Two hours after the doxycycline pulse, puromycin was added to terminate translation, and imaging was started 30 minutes after puromycin addition. All live-cell imaging was carried out at 37 °C. At the start of imaging, positions were selected based on the presence of SERP-expressing cells. Images were acquired over a period of 2-4 hours at a frame rate of 1 or 0.67 frames per minute.

### Data analysis

#### Flatfield correction

Flat-field processing was carried out using calibration images generated from a uniform fluorescent dye solution (4 µg/mL DyLight™ 488 NHS Ester for the 488-nm excitation) together with corresponding dark-current images. To generate the flat-field reference, 100 images were acquired from different stage positions and merged by calculating the median intensity at each pixel. Dark-field images were produced using identical camera settings but without laser light. A correction matrix was then generated by removing the dark-field contribution from the flat-field reference and scaling the resulting image to its average intensity. For all analyzed microscopy data, each frame was corrected by first subtracting the dark-field image and then dividing the result by the correction matrix.

#### Single-molecule tracking and fluorescence quantification

For tracking and measuring intensities of SERPs over time, we used the TransTrack software package developed previously^57^. To ensure that the full GFP spot was captured in the region of interest (ROI) used to track spots, we used a large, 8×8-pixel ROI, which was large enough to encompass the entire SERP spot. Photobleaching correction of intensity traces was performed as described previously^22^.

#### Calculation of degradation rates

To determine absolute proteasomal degradation rates from fluorescence intensity–time traces of degrading SERPs, the following calibration procedure was applied:

1. Fluorescence calibration: A reference protein containing exactly 24 SunTag epitopes was tethered to the plasma membrane via a CAAX motif and imaged under identical conditions in the same cell line. This allowed determination of the fluorescence intensity conferred by a single SunTag epitope.
2. Epitope number calculation: For each SERP, total fluorescence intensity was divided by the single-epitope intensity from (1) to calculate the number of SunTag epitopes per SERP.
3. Conversion to amino acids: The number of SunTag epitopes per SERP was next converted into the total size of a SERP in amino acids by multiplying SunTag epitope number from (2) by the average SunTag epitope spacing (in amino acids). The average SunTag epitope spacing was calculated from the known sequence of each socRNA (e.g., a socRNA encoding 5 SunTag epitopes over 200 amino acids yields an average spacing of 40 amino acids per epitope).
4. Degradation slope fitting: Fluorescence intensity-time traces of degrading SERPs were first converted to amino acid units using the above steps and then fit with a linear regression model to extract the slope of the degradation phase. As imaging was performed at 1-minute intervals, the resulting slope reflects the number of amino acids degraded per minute.
5. Conversion to per-second rate: Finally, the slope was divided by 60 to yield degradation speed in amino acids per second.

#### Calculation of plateau duration between the end of translation and the beginning of decay

To determine the time from ribosome quality control initiation until substrate degradation by the proteasome, we examined fluorescence intensity time-traces. GFP intensities initially increase, reflecting socRNA translation. At some point, a plateau is observed in the GFP intensity-time trace followed by a rapid decrease in intensity, reflecting ribosome stalling followed by activation of the ribosome quality control pathway and SERP degradation, respectively. To determine the time from ribosome quality control activation until SERP degradation, the plateau phase duration must thus be determined. (Fig. 5e,f and Supplementary Fig. 4d). To calculate this plateau phase duration, a sequential fitting strategy was applied, which served to define the moment when SERP synthesis was halted and the moment when decay was initiated. The time between these two moments reflects the plateau duration.

1. First fit to determine onset of the decay phase: The onset of signal decay was first defined by fitting a linear regression model from the last data point of the decay phase to the first data point that unambiguously belonged to the decay phase. Next, a residual plot of that linear regression to all data points of the trace (including data points of the signal increase or plateau phase) was plotted. This residual plot exhibited a clear bimodal distribution, separating data points belonging to the decay phase and data points that did not. This distribution was then used to demarcate the transition between data points belonging to the decay phase and decay points belonging to the signal increase and plateau phase.
2. Second fit to determine onset of plateau phase: To determine the moment of transition from signal increase to signal plateau, all data points before the transition to decay (1) were fit with a two-state linear regression model. The first segment of the fit was constrained to have a positive slope, reflecting continued synthesis, while the second was fixed at a slope of zero, reflecting a stable plateau. The intersection of these two fits was defined as the moment of onset of the plateau phase.
3. To calculate the duration of the plateau phase, the time between the moment of plateau onset (2) and the moment of decay onset (1) was calculated

#### Degradation kinetics of SERPs with sub-stoichiometric SunTag labeling

To assess whether antibody-based labeling of SunTag epitopes interferes with proteasomal degradation, we examined degradation kinetics in conditions with sub-stoichiometric decoration of SunTag epitopes. If the proteasome pauses upon encountering an antibody, reduced antibody labeling of the SERP should reduce the number of pauses and thus increase the average decay speed. Here we will explain how sub-stoichiometric labeling was induced and quantified. To achieve sub-stoichiometric epitope labeling, U2OS cells stably expressing TetR and ALFA–Nb–CAAX were transduced with a lentiviral construct encoding STAb–mStayGold under the control of a truncated SV40 promoter (dsV40), which is a weak promoter and yields low expression, generating a polyclonal population of cells with low and heterogeneous STAb expression levels. Very low levels of STAb-mStayGold in some cells resulted in sub-stoichiometric labeling of SunTag peptide epitopes.

To determine the precise labeling stoichiometry of SunTag peptides, cells were transfected with a let-7a–containing socRNA construct, and the rate of GFP increase during SERP synthesis was assessed. Since translation rates are not affected by nascent chain labeling, the number of SunTag peptides that are produced per unit of time is constant, irrespective of STAb-mStayGold expression. Thus, the rate of signal increase during SERP synthesis depends solely on the fraction of SunTag peptide epitopes that is labeled. Indeed, a positive correlation was observed between cytoplasmic STAb–mStayGold expression levels and the slope of fluorescence increase during SERP synthesis, consistent with variable SunTag peptide epitope labeling efficiency at variable STAb-mStayGold expression levels. Epitope occupancy across the STAb-mStayGold expression range was calculated by fitting the data in which total STAb-mStayGold expression levels were plotted against the rate of GFP increase of translated socRNAs to an exponential model that includes a plateau, which accounts for epitope saturation above which foci intensity cannot increase (Supplementary Fig. 1f).

#### Degradation rate of a SERP with a mNeonGreen repeat sequence

U2OS cells stably expressing TetR and ALFA–Nb–CAAX were transfected with a socRNA construct encoding an N-terminal 5×ALFA-tag array followed by mNeonGreen, the solubility tag GB1 and a let-7a target site within the socRNA coding sequence. Time-lapse imaging and SERP tracking were performed as described in ‘*Single-molecule tracking and fluorescence quantification*’. To calculate absolute proteasomal degradation speed, fluorescence decay slopes were compared to the slope of fluorescence increase per ribosome, measured in the same experiment. Since we know that the translation rate on socRNAs in these cells is 2.6 aa/s (Supplementary Fig. 1g), the decay rate could be calculated too; The slope of the decay phase was divided by the slope of SERP synthesis phase and multiplied by 2.6 to get the decay rate in aa/s.

#### Calculating the unfolding time of mDHFR during decay

To calculate the time required for the proteasome to unfold DHFR in the presence or absence of MTX, we first determined the average degradation rate on a control sequence lacking DHFR (Fig. 2e). Using this rate, we then calculated the expected degradation time of a control sequence matched in length to DHFR. To achieve this, we first calculated the average decay rate per amino acid for the control reporter lacking the DHFR sequence and then calculated the expected decay time of DHFR based on the number of amino acids. The unfolding time for DHFR was then determined by subtracting the expected time for degrading the DHFR containing SERP from the observed time.

#### Pause detection using Hidden Markov modeling

To detect pauses during proteasomal degradation, we analyzed signal decay intensity-time traces of individual degraded SERPs. Fluorescence traces were first smoothed using a two-point moving average, and the first derivative was calculated of the curve, representing intensity changes between consecutive time points. Pause detection was performed using a two-state Hidden Markov Model (HMM) approach based on the vbFRET algorithm^58^ to detect a maximum of 2 states per trace. These states reflect either a decay state (negative value for the intensity-time trace derivative) or a pause state (derivative value of close to zero).

To estimate the rate of false-positive pause detection caused by technical noise in the data, we generated a negative control dataset consisting of plateau traces acquired from MZB-treated cells. These plateau traces retain experimental noise but lack true decay phases due to proteasomal inhibition. These control traces underwent a linear transformation such that they had identical average decay slopes as the decay traces in the dataset. Also, the duration of control traces was matched to decay traces. The same HMM algorithm was applied to determine state values for each decay event in both datasets. While most traces exhibited negative state values, which correspond to steady fluorescence decay (i.e. SERP degradation), some traces displayed transient shifts to higher state values, reflecting proteasome pausing events. The highest state values per trace from the control data set were then used to define a detection threshold to call a bona fide pause. Traces were classified as undergoing a pause if their highest state value exceeded the cutoff derived from the control data set (mean + 2 s.d. of the control distribution).

Pausing events occurred at a low frequency, resulting in low fraction of traces exhibiting pauses in each individual experiment. This low number of pause events led to highly variable estimates of pause frequency between experiments, making it difficult to determine whether pause frequency in decay traces was statistically different from the observed pause frequency in control. To overcome this problem and to obtain a more robust estimate of uncertainty in the fraction of decay traces exhibiting a pause, decay traces from all experiments were therefore pooled and a resampling (bootstrapping) approach was applied.

Bootstrap datasets were generated by randomly sampling decay traces with replacement from the pooled dataset, while maintaining the original number of traces per condition. Equivalent bootstrap datasets were generated for control traces. For each bootstrap dataset, pause detection was performed using the threshold-based classification described above, and the fraction of traces containing a pause was calculated. This procedure was repeated 1,000 times, yielding distributions of pause fractions for both decay and control conditions. Error bars represent the standard deviation of these bootstrap distributions (Fig. 1o).

#### Quantification of the fraction of SERPs that is degraded

For the quantification of the fraction of SERPs that underwent degradation in the C-to-N degradation assay (Fig. 3i,j), SERPs were only included if they could be tracked for at least 40 minutes following plateau onset (which reflects the end of translation) to assess whether degradation occurred.

In the N-to-C degradation assay, degradation typically occurred within the first 10,000 translated codons (∼1 hour; Supplementary Fig. 2d). SERPs were marked as non-degraded if degradation did not occur within synthesis of 15,000 codons.

#### Quantification of degradation kinetics

To quantify degradation kinetics in internal proteasome recruitment experiments (Fig. 5j,l and Supplementary Fig. 4f,g), U2OS cells expressing SERPs encoding an internal dTAG or poly-leucine degron were analyzed. SERP intensity and number was determined using the *spot_Counter* plug-in in ImageJ with an 8-pixel box size. For each time-point and each field of view, total spot intensities were calculated by multiplying the number of spots by their average intensity. Values were normalized either to the start of imaging (Fig. 5l) or to the time of dTAG-v1 addition (Fig. 5j and Supplementary Fig. 5b,c).

#### Calculating the number of amino acids degraded prior to fragmentation for substrate-internal proteasome engagement

To quantify the number of amino acids degraded prior to the moment of SERP fragmentation under 5 nM dTAG-v1 treatment, fluorescence intensity-time traces were first aligned to the time point when visible foci splitting occurred. Because fragmentation of a single SERP becomes detectable only after the two protein fragments have diffused sufficiently apart to be resolved as distinct diffraction-limited foci, we corrected for the time lag between the actual fragmentation event and its visual detection. We have previously determined the time between splitting of two GFP foci and the visible detection of this splitting under similar conditions to be 2.0 minutes^22^. We therefore set the moment of foci fragmentation to two minutes before fragmentation was first visible (Fig. 5n).

#### Outcome of proteasome-ribosome collisions

To distinguish between possible outcomes of proteasome–ribosome collisions, we compared the slope of intensity-time traces obtained before and after a degradation event, or before and after the addition of MZB. The slope of the intensity–time trace directly reports the number of ribosomes translating a socRNA, as each translating ribosome produces SunTag peptides that contribute to the fluorescent signal (Supplementary Fig. 3b). Notably, in ‘Model 2’ (Fig. 4a), the ribosome is producing SunTag peptides, but they are immediately degraded by the trailing proteasome and are therefore not fluorescently labeled and are not contributing to the slope.

To test whether proteasomes release the substrate upon collision with the ribosome (Model 1 versus Models 2 or 3), we compared the rate of spot intensity increase before the decay event with the rate of spot intensity increase after the decay event (Fig. 4e). To ensure that only events corresponding to translation by two or more ribosomes during the 1^st^ signal increase phase were included in the analysis, we applied a cutoff where the fluorescence slope had to exceed 1.75× the median slope measured for single translating ribosomes. Additionally, to exclude cases where proteasomes dissociated from the nascent protein chain before reaching the ribosome, the duration of the decay phase was required to be at least half as long as the 1^st^ increase phase. Finally, both the first and second increase phases were required to contain at least 10 data points, reducing noise in slope measurements. MZB was added 90 minutes after the start of imaging, enabling discrimination between Model 2 (proteasome trailing) and Model 3 (ribosome dissociation) based on the resulting changes in slope (Fig. 4h and Supplementary Fig. 3c).

#### Calculation of proteasome-ribosome collisions on endogenous mRNAs

To estimate the furthest position along a coding sequence (CDS) on an endogenous mRNA at which a co-translationally engaged proteasome can still catch up with a translating ribosome (i.e. before the ribosome reaches the stop codon), we modelled a scenario in which the ribosome and proteasome move at constant velocities. We used our experimentally determined rates of 2.6 aa/s for ribosomal translocation and 7.8 aa/s for proteasomal N-to-C degradation (Fig. 3d and Supplementary Fig. 1g).

We define *L* as the length of the CDS (in amino acids), measured from the start codon to the stop codon. We define *p*_eng_ as the position of the ribosome along the CDS at the moment when the proteasome engages the nascent polypeptide at its N-terminus. At this moment, the ribosome has already translated *p*_eng_amino acids, and *L* − *p*_eng_amino acids remain to be translated.

After proteasome engagement, the ribosome continues translation elongation at a velocity 𝑣_*r*_, while the proteasome degrades the nascent chain from the N-terminus toward the C-terminus at a velocity 𝑣_*p*_. The time remaining until translation termination by the ribosome (*t*_term_) is therefore:

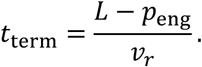

During this remaining time, the proteasome degrades the nascent chain and closes the distance to the ribosome while the ribosome continues to translate towards the end of the CDS until it reaches the stop codon at position L. Consequently, the distance between the proteasome and the ribosome decreases at the relative velocity 𝑣_*p*_ − 𝑣_*r*_. The time required for the proteasome to reach the ribosome (*t*_reach_) is thus:

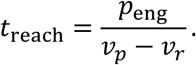

The proteasome can reach the ribosome before translation termination when the time required to reach the ribosome is shorter than the time remaining until termination, that is,

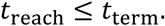

Substituting the expressions above gives

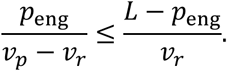

This condition can be rearranged to yield the equivalent expression

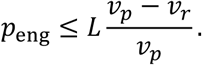

Solving this inequality yields a maximal ribosome position *p*_eng_ at which proteasome recruitment and degradation towards the C-terminus can still result in a proteasome-ribosome collision before the moment of translation termination. Using the experimentally measured values 𝑣_*p*_ = 7.8 amino acids per second for protein degradation rate and 𝑣_*r*_ = 2.6 amino acids per second for the translation elongation rate, this maximal position corresponds to approximately 67% of the CDS length. Thus, under these assumptions, a co-translationally recruited proteasome can catch up with and collide with a translating ribosome before translation termination only if engagement occurs before 67% of the CDS has been translated.

## Statistical analysis

Statistical analyses were conducted in GraphPad Prism (version 9.4.1). Details of statistical tests and error bars are provided in figure legends or directly within the figures.

## Data availability

Requests for reagents, resources and code should be directed to and will be fulfilled by the Lead Contact, Marvin Tanenbaum. Plasmids used in this study will be deposited and made available in Addgene. A selection of raw imaging from this study has been made available on Mendeley.

## Acknowledgements

We thank members of the Tanenbaum lab for helpful discussions, S. Yang for providing code and support with Hidden Markov modeling, J. Schokolowski, M. Müller and M. Baars for help with data visualization. M.F.M. and M.E.T. were supported by the Oncode Institute, which is partly funded by the Dutch Cancer Society (KWF). M.E.T. acknowledges funding from the VIDI (NWO/016.VIDI.189.005). D.H was supported by Boehringer Ingelheim.

---

**Supplementary Fig. 1.**
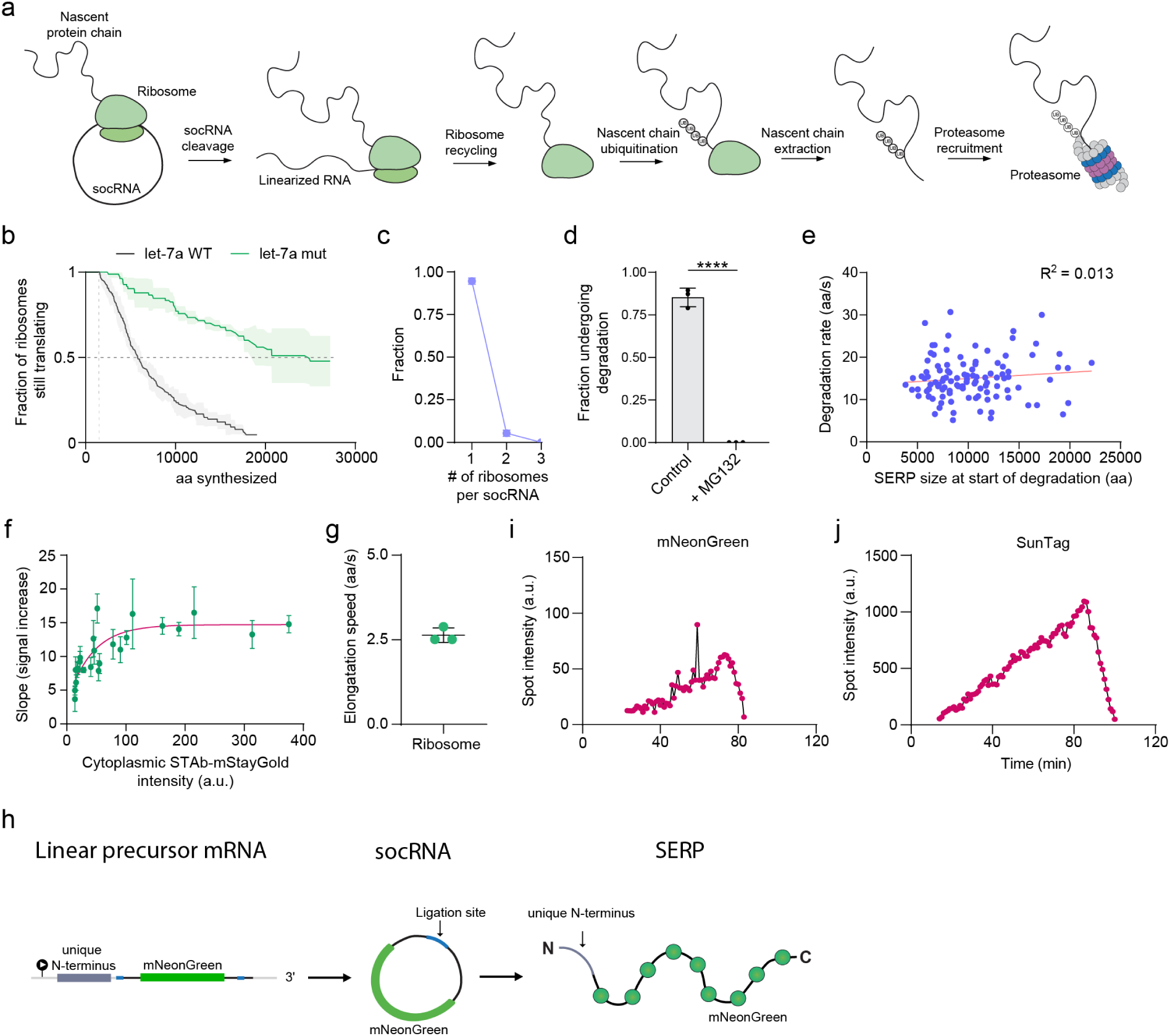
Degradation kinetics of single proteasomes in living cells. **a,** Schematic of the ribosome quality control pathway co-opted to induce SERP degradation. Cleavage of the socRNA results in a terminally stalled ribosome that is recognized by quality control factors, triggering ribosome splitting and dissociation from the mRNA. The resulting 60S–nascent chain complex is then recognized and the nascent chain ubiquitinated, leading to extraction of the nascent protein and its subsequent delivery to the proteasome for degradation. **b,** Kaplan-Meier survival curves showing the number of codons translated per ribosome on the indicated socRNA before translation elongation is aborted. Solid line and shaded area represent mean ± s.d (n = 126 (wt) and 90 (mut) traces from 3 independent repeats). **c,** Distribution of the number of translating ribosomes and associated SERPs per socRNA. Horizontal line and error bars represent mean ± s.d (n = 124 socRNAs from 2 independent repeats). **d,** Fraction of SERPs undergoing degradation in untreated cells and in cells treated with the proteasome inhibitor MG132. Horizontal line and error bars represent mean ± s.d. Unpaired Student’s t-test was used for statistical analysis. **** denotes p < 0.0001 (n = 136 traces (control) and n = 54 traces (+MG132) from 3 independent repeats). **e,** Relationship between SERP size and the rate of degradation. The red line indicates a linear fit to the data, and the coefficient of determination (R²) is shown (n = 113 traces from 4 independent repeats). **f,** Relationship between cytoplasmic STAb-mStayGold expression levels and the slopes of the signal increase phase (SERP synthesis rate by ribosomes). Since translation rates are independent of STAb-mStayGold expression, the lower slope at lower STAb-mStayGold expression reflects sub-stoichiometric labeling of newly-synthesized SunTag epitopes. Each dot and set of error bars represents the average ± s.d of all the events in a single cell. Red line indicates exponential plateau fit to the data (n = 151 SERPs over 24 cells). **g,** Translation elongation speed of single translating ribosomes. Ribosome speed was measured for socRNA shown in Fig. 1c. Horizontal line and error bars represent mean ± s.d of medians (n = 100 traces from 3 independent repeats). **h,** Schematic illustrating the strategy to generate SERPs with a unique N-terminus followed by a repeating array of mNeonGreen proteins. A unique N-terminal protein sequence is encoded in the linear precursor mRNA, which is translated only once by the ribosome before RNA circularization occurs. The mNeonGreen sequence is encoded in the repeat unit of the mRNA, the part that is excised and circularized. **i,** Representative fluorescence intensity trace showing SERP synthesis (signal increase) followed by SERP degradation (signal decay). SERPs were fluorescently labeled using mNeonGreen encoded in the socRNA, as shown in (h). **j,** Representative fluorescence intensity trace showing SERP synthesis (signal increase) followed by SERP degradation (signal decay). SERPs were fluorescently labeled using SunTag epitope arrays encoded in the socRNA, as shown in Fig. 1b. Different imaging settings were used in (i) and (j); therefore, fluorescence intensities cannot be directly compared between the two panels. However, SERPs labeled using the SunTag-based approach were brighter resulting in lower observed noise in intensity-time traces.

**Supplementary Fig. 2.**
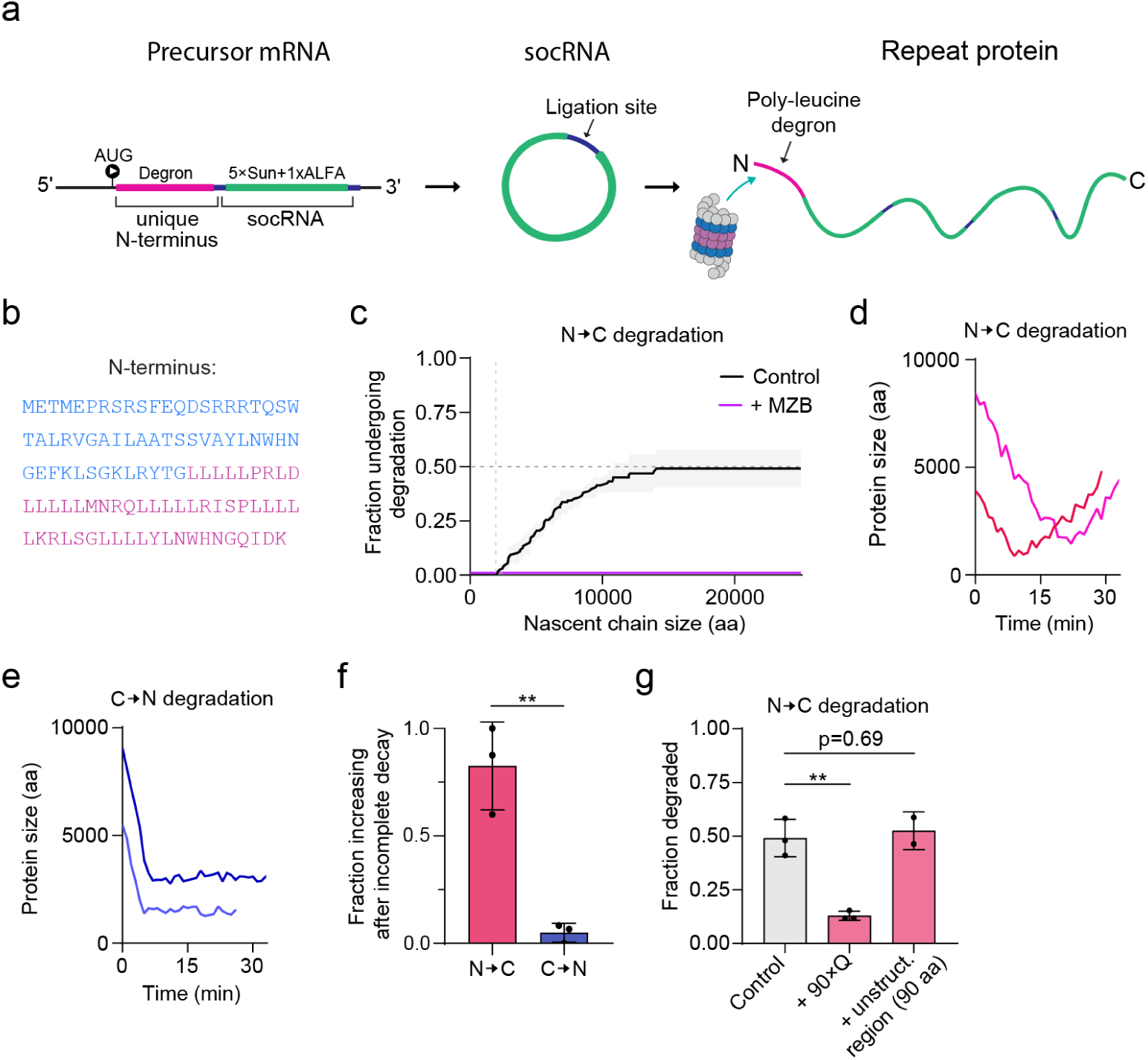
Effect of proteasome directionality on degradation. **a,** Schematic of the strategy used to recruit proteasomes to the SERP N-terminus. A unique N-terminal poly-leucine degron is encoded in the linear precursor mRNA and is translated only once by the ribosome before RNA circularization occurs. The 5×SunTag and 1×ALFA-tag are encoded within the repeat region that is excised and circularized to form the socRNA. **b,** Sequence of the N-terminal degron. The region highlighted in blue denotes an unstructured proteasome initiation region derived from cytochrome b2. The region highlighted in pink denotes the poly-leucine-rich degron sequence. The unique N-terminus was encoded upstream of the SERP repeat unit, as illustrated in (a). **c,** Cumulative incidence curve showing the fraction of SERPs subjected to N-to-C degradation. Treatment of cells with MZB prevents degradation. Line and shaded region indicate mean ± s.d (n = 130 traces (control) and n = 37 traces (+MZB) from 3 independent repeats). **d,** Representative intensity-time traces of SERPs undergoing incomplete N-to-C degradation. Incomplete degradation results in resumption of signal increase as the C-terminus continues to be extended by the translating ribosome, as N-to-C degradation occurs co-translationally. **e,** Representative intensity-time traces of SERPs undergoing incomplete C-to-N degradation. Incomplete degradation results in a sustained signal plateau, as C-to-N degradation occurs post-translationally. **f,** The fraction of decay events showing intensity increase following incomplete decay for either N-to-C or C-to-N degradation assays. Horizontal line and error bars represent mean ± s.d (n = 22 traces (N-to-C) and n = 58 traces (C-to-N) from 3 independent repeats). **g,** Fraction of indicated SERPs undergoing degradation in the N-to-C decay assay. Horizontal line and error bars represent mean ± s.d (n = 130 traces (control), n = 58 traces (+90xQ) and n = 66 traces (+unstructured region 90aa) from 2-3 independent repeats). Control data is replotted from Fig. 3c. **(f, g),** Unpaired Student’s t-test was used for statistical analysis. * and ** denote p < 0.05 and 0.01, respectively.

**Supplementary Fig. 3.**
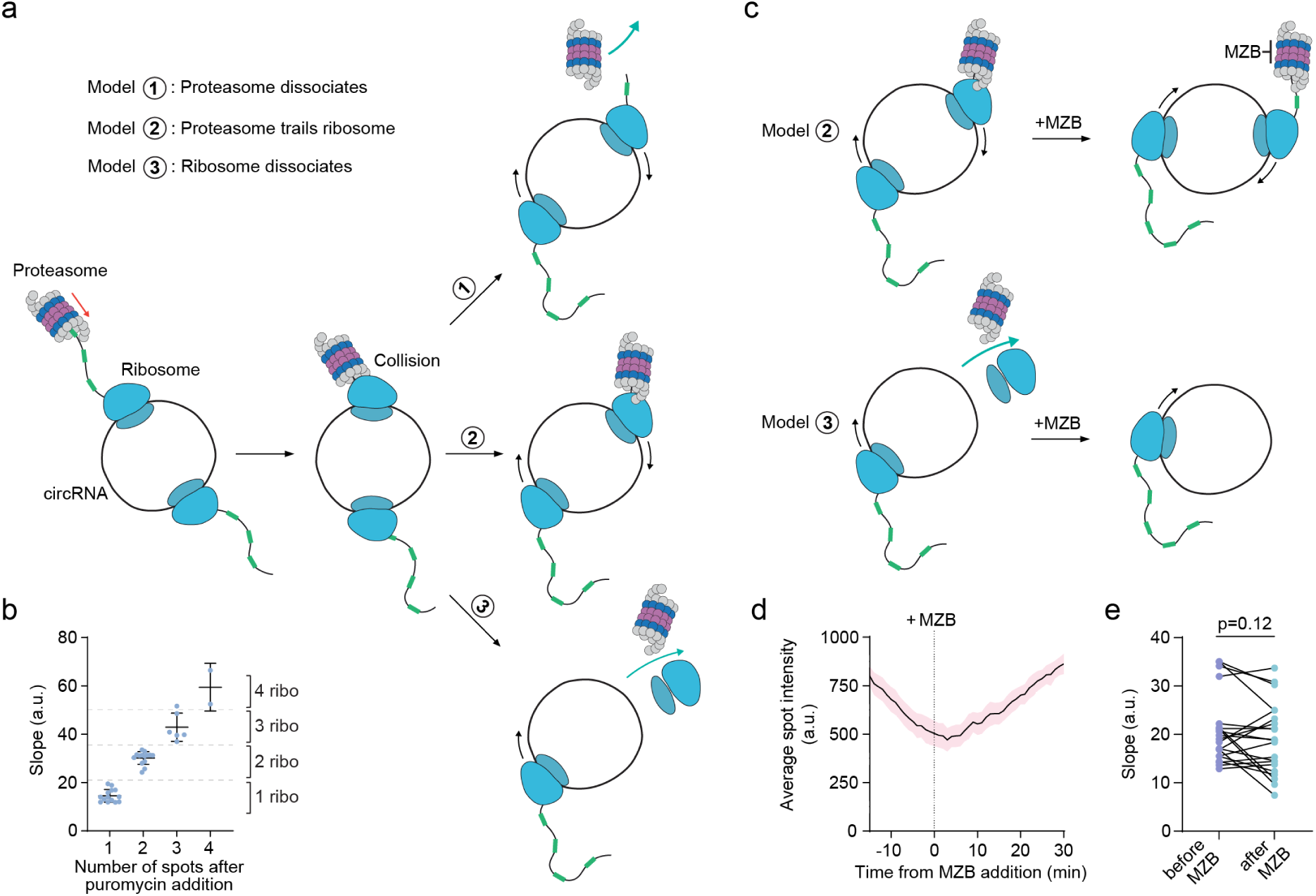
Proteasome-ribosome collisions promote ribosome dissociation. **a,** Cartoon illustrating different outcomes of proteasome-ribosome collisions on a socRNA translated by 2 ribosomes. A single proteasome co-translationally degrades one of the two nascent chains. Upon collision with the ribosome, the proteasome either dissociates allowing the ribosome to continue translation (Model 1), continues to trail the translating ribosome (Model 2), or induces ribosome dissociation from the mRNA (Model 3). **b,** Relationship between slope of the signal increase phase and number of ribosomes translating the socRNA, as assessed by the number of GFP foci upon puromycin treatment. Horizontal line and error bars represent mean ± s.d (n = 36 traces from 1 repeat). **c,** Cartoon illustrating possible outcomes of proteasome inhibition with MZB following proteasome collision with one of two ribosomes translating a socRNA (as shown in **(a)**). In Model 2 (proteasome trailing), MZB inhibits continued decay of the newly synthesized nascent chain, resulting in an increase in GFP signal accumulation. In Model 3, MZB has no effect on GFP accumulation rates, as the ribosome has been displaced from the socRNA. **d,** Average SERP fluorescence intensity during degradation, aligned in time to the moment of MZB addition. Dashed vertical line indicates the moment of MZB treatment. Solid line and shaded region indicates mean ± s.d (n = 9 traces from 3 independent repeats). **e,** Comparison of the slopes of the 2^nd^ increase phase before and after MZB addition, using traces in which the 1^st^ increase phase reflected translation by 2 or more ribosomes. Paired Student’s t-test was used for statistical analysis. Each set of dots reflects a single SERP. Lines connect the same SERP before and after MZB treatment (n = 21 traces from 3 independent repeats).

**Supplementary Fig. 4.**
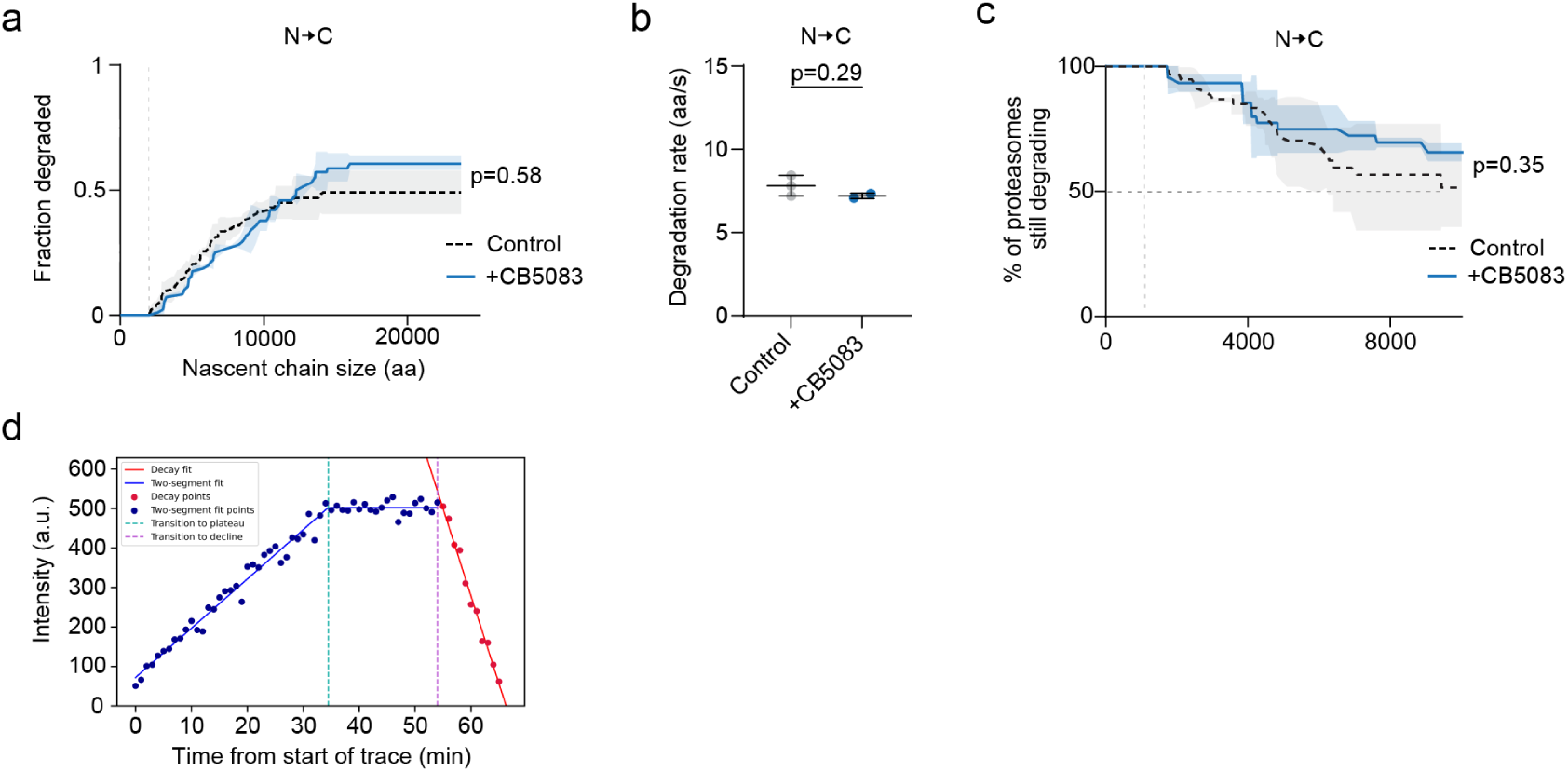
p97 is dispensable for N-to-C proteasomal degradation. **a,** Cumulative incidence curve showing fraction of SERPs subjected to N-to-C degradation in the presence or absence p97 inhibitor CB5083. Control data is replotted from Supplementary Fig. 2c for comparison (n = 130 traces (control) and n = 70 traces (+CB5083) from 2-3 independent repeats). **b,** Proteasomal degradation rate for N-to-C degradation in the presence or absence of CB5083. Each data point represents the median from an independent experiment. Horizontal line and error bars represent mean ± s.d of medians (n = 62 traces (control) and n = 36 traces (+CB5083) from 3 independent repeats). Unpaired Student’s t-test was used for statistical analysis. **c,** Kaplan-Meier survival curves showing proteasomal processivity for N-to-C degradation in the presence or absence of CB5083. Line and shaded region indicate mean ± s.d (n = 62 traces (control) and n = 36 traces (+CB5083) from 2-3 independent repeats). Control data is replotted from Fig. 3e. **d,** Data fitting approach for the trace shown in Fig. 5e to identify the onset of ribosome stalling and the onset of protein degradation. **(a,c),** Log-rank (Mantel-Cox) test was used for statistical analysis.

**Supplementary Fig. 5.**
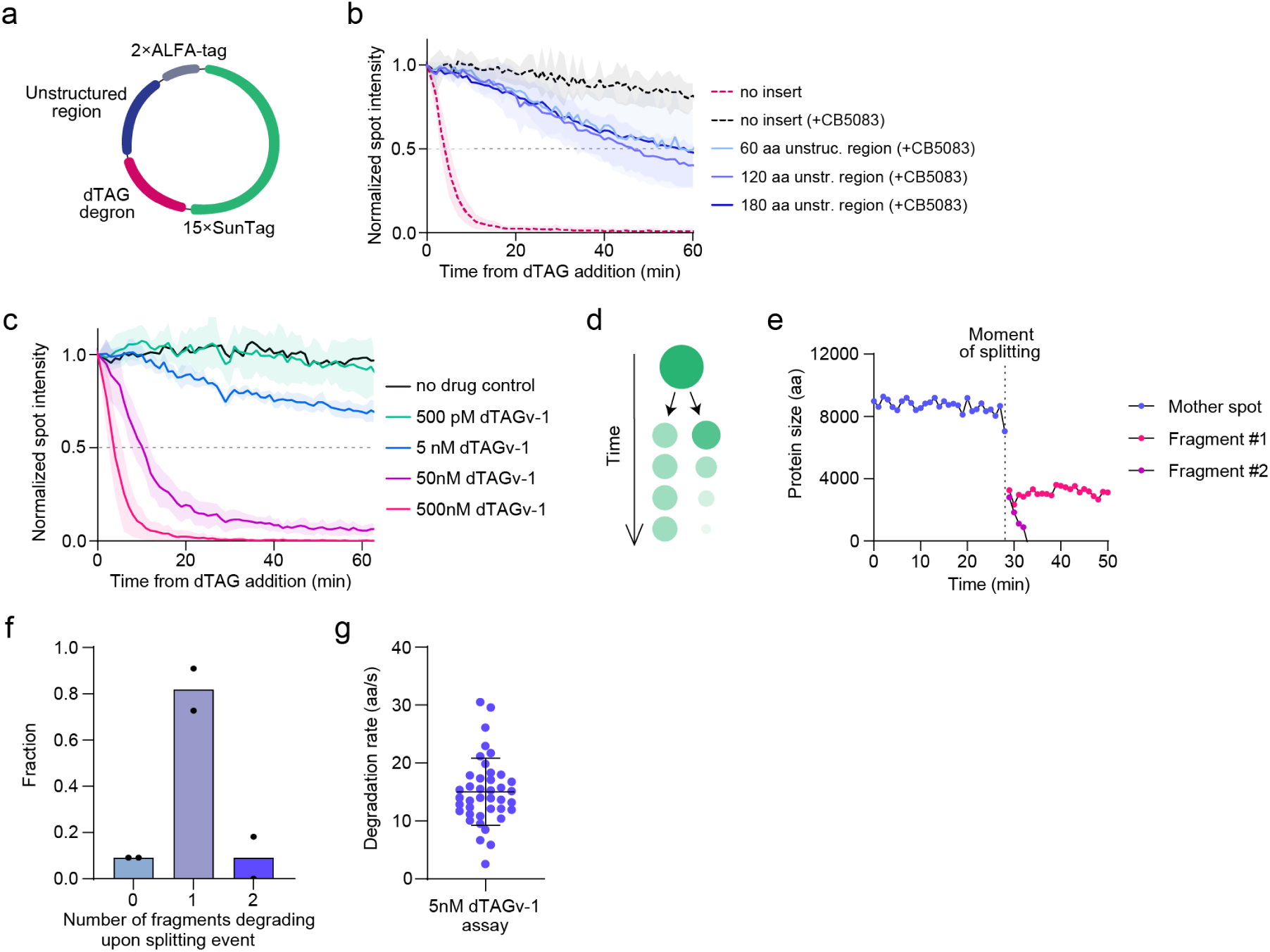
Analysis of substrate-internal proteasomal degradation. **a,** Schematic of the socRNA used for substrate-internal proteasome recruitment experiments. The socRNA encodes 15× SunTag repeats, 2× ALFA-tag repeats, a single dTAG-inducible degron and an unstructured region of 60, 120 or 180 amino acids in size. **b,** Quantification of SERP degradation kinetics following addition of 500 nM dTAGv-1. Lines and shaded region indicate mean ± s.d (n = 8 fields of view (no insert control), n = 7 fields of view (no insert +CB5083), n = 7 fields of view (60 aa unstructured region +CB5083), n = 6 fields of view (120 aa unstructured region +CB5083) and n = 8 fields of view (180 aa unstructured region +CB5083) from 2-3 independent repeats). Dotted lines representing control SERP are replotted from Fig. 5j. **c,** Quantification of SERP degradation kinetics following addition of varying concentrations of dTAGv-1. SERPs were produced from socRNA shown in Fig. 5h. Lines and shaded region indicate mean ± s.d (n = 7 fields of view (no drug control), n = 6 fields of view (500 pM dTAGv-1), n = 7 fields of view (5 nM dTAGv-1), n = 6 fields of view (50 nM dTAGv-1) and n = 6 fields of view (500 nM dTAGv-1) from 2 independent repeats). **d,** Schematic depicting fragmentation of SERP as a consequence of substrate-internal proteasome recruitment. Upon fragmentation, one of the two SERP fragments is degraded, while the other fragment remains stable over time. **e,** Representative intensity time-trace of a SERP undergoing fragmentation and degradation in the presence of 5 nM dTAGv-1. One of the two SERP fragments is completely degraded while the other SERP fragment remains stable over time. **f,** Quantification of number of SERP fragments that underwent degradation after fragmentation into two fragments. Bars represent the average of 2 independent repeats, individual experiments are shown as dots (n = 22 traces from 2 independent repeats). **g,** Rate of SERP degradation induced by 5 nM dTAGv-1. Data points represent individual degradation events. Horizontal line and error bars represent mean ± s.d (n = 40 traces from 2 independent repeats).

## Supplementary information

**Supplementary Table 1.** Number of experimental repeats and details on construct sequences used in this study.

**Supplementary Video 1 | Live-cell imaging of protein degradation by single proteasomes.**

Representative movie showing three individual SERPs undergoing proteasomal degradation. SERPs were produced from the socRNA shown in Fig. 1c. Degradation in C-to-N directionality was induced through socRNA cleavage as shown in Supplementary Fig. 1a. SERPs were expressed in U2OS cells stably expressing STAb-mStayGold, ALFANb-CAAX, and TetR. Images were acquired at 60 s intervals. Scale bar, 1 µm.

**Supplementary Video 2 | Visualization of proteasomal processivity at the single-molecule level.**

Representative movie showing three individual SERPs undergoing proteasomal degradation. For one out of three SERPs, signal decay is incomplete, indicating that the substrate dissociated from the proteasome before being completely degraded. SERPs were produced from the socRNA shown in Fig. 1c. SERPs were expressed in U2OS cells stably expressing STAb-mStayGold, ALFANb-CAAX, and TetR. Images were acquired at 60 s intervals. Scale bar, 1 µm.

**Supplementary Video 3 | p97 inhibition blocks substrate-internal degradation.**

Representative movies showing rapid SERP degradation and inhibition of degradation by p97 inhibition. SERPs encoded 15×SunTag epitopes, 2× ALFA-tag epitopes and 1×dTAG degron per repeat unit. Cells were pre-treated with the p97 inhibitor CB5083 (5 µM; right) for 10 minutes or left untreated (left). The dTAG ligand dTAGv-1 was added to both conditions at t = 5 min. SERPs were expressed in U2OS cells stably expressing STAb-mStayGold, ALFANb-CAAX, and tetR. Images were acquired at 60 s intervals. Scale bar, 10 µm.

